# Linker histones regulate fine-scale chromatin organization and modulate developmental decisions in Arabidopsis

**DOI:** 10.1101/458364

**Authors:** Kinga Rutowicz, Maciej Lirski, Benoît Mermaz, Jasmin Schubert, Gianluca Teano, Imen Mestiri, Magdalena A. Kroteń, Tohnyui Ndinyanka Fabrice, Simon Fritz, Stefan Grob, Christoph Ringli, Lusik Cherkezyan, Fredy Barneche, Andrzej Jerzmanowski, Célia Baroux

**Affiliations:** Institute of Plant and Microbial Biology, Zürich-Basel Plant Science Center, University of Zürich, Switzerland; Institute of Biochemistry and Biophysics, Polish Academy of Sciences, Pawinskiego 5a, 02-106 Warsaw, Warsaw, Poland; Institut de Biologie de l’Ecole Normale Supérieure (IBENS), Ecole normale supérieure, CNRS, INSERM, PSL Research University, 75005 Paris, France; Département de Biologie, IBENS, Ecole Normale Supérieure, CNRS, INSERM, PSL Research University, 46 rue d’Ulm, F-75005, Paris, France; College of Inter-Faculty Individual Studies in Mathematics and Natural Sciences, University of Warsaw, 02-089 Warsaw, Poland; Department of Biomedical Engineering, Northwestern University, Evanston, Illinois 60208, USA; Faculty of Biology, University of Warsaw, Pawinskiego 5a, 02-106 Warsaw, Warsaw, Poland

**Keywords:** Linker histones, H1, chromatin, heterochromatin, histone methylation, developmental transitions

## Abstract

Chromatin in eukaryotes provides a tunable platform to control gene expression and convey an epigenetic memory throughout cell divisions. H1 linker histones are abundant components with an intrinsic potential in influencing chromatin structure and function. We detail the impact of H1 depletion in Arabidopsis on fine-scale chromatin organization, transcription and development. While required for chromocenter assembly, H1s are dispensable for transposable element (TE) silencing and peripheral positioning of heterochromatin. In euchromatin, H1 regulates nucleosome density, mobility, and regular distribution of nanoscale chromatin domains. While necessary to maintain epigenetic patterns, H1 only moderately affects transcription. Its depletion is associated with failures in transitional fate changes such as lateral root initiation, root hair production, stomata patterning but also flowering and dormancy regulation. Therefore, Arabidopsis H1 variants are chromatin architects mediating nano‐ and microscale levels-of-organization operating downstream of epigenetic and transcriptional establishment processes and contribute to epigenetic reorientations in developmental transitions.

## Introduction

Linker histones (H1) are one of the major components of plant and animal chromatin. H1 (referring to the entire variants family) appeared early during evolution, with lysine-rich proto-linker histones found in the most ancestral eukaryotes (Kasinsky, Lewis et al. 2001). In contrast to the core nucleosomal constituents, however, H1 is the most divergent class of histones (Kasinsky, Lewis et al. 2001). H1 typically possess a conserved tripartite structure composed of a short and flexible N-terminal tail, a structured globular domain (GH1) which interacts with a nucleosome dyad and a structurally disordered, lysine-rich (highly basic) C-terminal tail. The C-terminal tail, which varies in length and composition among variants and organisms, interacts with internucleosomal linker DNA and draws adjacent nucleosomes together thus conferring the chromatin compaction potential of H1 variants (Zhou, Feng et al. 2013, Bednar, Garcia-Saez et al. 2017). Several variants can co-exist in one cell playing then redundant and specific roles in chromatin structure and functions (reviewed in (Fyodorov, Zhou et al. 2018). H1 proteins constitute a highly mobile fraction of the chromatin and their apparent constitutive presence results from a steady-state level of dynamic binding (Bustin, Catez et al. 2005). H1 variants differ in DNA and nucleosome binding properties, regulating both chromatin compaction at mitosis as well as nucleosomal spacing at interphase (reviewed in (Hergeth and Schneider 2015)). The structural role of H1 in chromatin organization has an influence on several genome functions such as gene regulation, DNA replication, chromosome segregation and DNA repair (reviewed in (Almeida, Fernandez-Justel et al. 2018, Fyodorov, Zhou et al. 2018)). Yet, the functional impact of H1 depletion shows a large variability depending on H1 variants and organisms. While H1 seems dispensable in *Tetrahymena thermophila*, yeast, and the fungus *Ascobolus immersus* (Shen, Yu et al. 1995, Ushinsky, Bussey et al. 1997, Patterton, Landel et al. 1998, Ausio 2000), its loss-of-function has a variable impact in higher organisms, ranging from developmental alterations in *Caenorhabditis elegans* and the flowering plant *Arabidopsis thaliana* (Jedrusik and Schulze 2001, Wierzbicki and Jerzmanowski 2005) to early lethality in mouse and Drosophila (Fan, Nikitina et al. 2005, Lu, Wontakal et al. 2009). Generally, H1 has been implicated in the control of genetic programs during development and differentiation (Kasinsky, Lewis et al. 2001, Hergeth and Schneider 2015, Pan and Fan 2016). Yet, H1 moderately impacts on global gene expression in mammalian cell cultures (Fan, Nikitina et al. 2005, Sancho, Diani et al. 2008, Zhang, Cooke et al. 2012, Geeven, Zhu et al. 2015) still affecting the expression of pluripotency genes (Zhang, Cooke et al. 2012). This is in line with the implication of H1 in regulating nucleosomal density and RNA Polymerase II accessibility in pluripotent cells (Christophorou, Castelo-Branco et al. 2014, Ricci, Manzo et al. 2015). H1 has also been shown to have a role in controlling epigenetic marks such as DNA methylation (Fan, Nikitina et al. 2005, Yang, Kim et al. 2013, Seymour, Ji et al. 2016) and histone H3 methylation (Lu, Wontakal et al. 2013, Yang, Kim et al. 2013, Geeven, Zhu et al. 2015). The intrinsic role of H1 on chromatin organization and yet moderate impact of its depletion on cell viability creates an apparent paradox that was early recognized (Bustin, Catez et al. 2005).

The plant kingdom possesses H1 variants that can be traced to earliest land plants (Kotlinski, Knizewski et al. 2017). The flowering plant Arabidopsis possesses three canonical H1 variants (Wierzbicki and Jerzmanowski 2005, Kotlinski, Knizewski et al. 2017). Two of them, H1.1 and H1.2 are canonical variants expressed throughout the plant except in cells of the reproductive lineage (She, Grimanelli et al. 2013, She and Baroux 2015, Ingouff, Selles et al. 2017). H1.3 is a stress-inducible variant contributing to physiological adaptation (Rutowicz, Puzio et al. 2015). Previous studies based on different reverse genetics approaches reported a variable impact of H1 depletion. RNAi-based downregulation of all three variants induces severe developmental aberration and sterility (Wierzbicki and Jerzmanowski 2005). By contrast, a stepwise introgression of insertional (T-DNA) genetic lesions generated viable plants, either double (*h1.1h1.2*, (Zemach, Kim et al. 2013)) or triple (*h1.1h1.2h1.3*, (She, Grimanelli et al. 2013, Rutowicz, Puzio et al. 2015) mutant lines. The triple mutant (thereafter called *3h1*) shows no detectable levels of H1 protein in immunostaining and immunoblot (She, Grimanelli et al. 2013), yet the plants are viable and do not exhibit dramatic morphological alterations. Possibly, and as already suggested for other organisms and earlier for Arabidopsis, the lack of H1 variants may be partially compensated (Jerzmanowski, Przewtoka et al. 2000, Bustin, Catez et al. 2005) for example by HMG-related proteins that are abundantly present in plant cells (Launholt, Gronlund et al. 2007).

H1 variants (thereafter referred collectively to as H1, for simplicity) are distributed across the genome, spanning both heterochromatin and euchromatin chromosomal domains (Rutowicz, Puzio et al. 2015).H1.1 and H1.2 variants are enriched at the 3’ and 5’ ends of TEs and over gene bodies anti-correlating with gene expression and H3K4me3 levels (Rutowicz, Puzio et al. 2015). Like in animals, Arabidopsis H1s have been recognized to have a dramatic impact on DNA methylation, altering patterns primarily but not exclusively in heterochromatin, and affecting all sequence contexts (Wierzbicki and Jerzmanowski 2005, Rea, Zheng et al. 2012, Zemach, Kim et al. 2013). However, the specific role of H1 on chromatin organization and function in plants remains elusive. This prompted us to investigate the detailed structure, composition and organization of H1-depleted chromatin in somatic plant cells. We found that H1 has distinct roles in heterochromatin and euchromatin organization at the microscopic and nanoscale level. Notably, H1 is necessary to maintain heterochromatin organization but dispensable for peripheral localization and epigenetic silencing of heterochromatin and has only a moderate influence on nucleosomal density. In euchromatin, H1 depletion results in a quantifiable dispersion of chromatin density patterns at the ultrastructural level corresponding to a loss of regularity in the distribution of locally dense chromatin regions (formerly called nucleosomal clutches in mammalian cells, (Ricci, Manzo et al. 2015). This correlates with an overall reduced nucleosome repeat length and altered nucleosomal occupancy which blurs the distinction between transcriptional‐ and epigenetic-dependent states of chromatin (Roudier, Ahmed et al. 2011, Sequeira-Mendes, Araguez et al. 2014). At the same time, H1 depletion induces hyperacetylation of the chromatin, and reduction of H3 methylation (but not H3 levels itself).

Our study thus uncovers a function for H1 in non-heterochromatic regions, which was so far overlooked. Yet, and reminiscent to findings in mammalian cells, H1 depletion has a moderate impact on gene expression, at least in standard plant growth conditions. Thus, our analysis reinforces the idea that H1-mediated, large-scale chromatin organization is dispensable for basic cellular functions and plant growth. Nevertheless, the observation of mild, yet specific phenotypes altering flowering transition, seed dormancy relief, lateral root formation, stomatal spacing, and competence to form callus under inducing conditions suggests a role for H1 in providing robustness during developmental transitions. We propose a model where H1-mediated chromatin organization, operating at the nanoscopic and nuclear scale level, facilitates transcriptional reprogramming under developmental cues.

## Results

### H1 variants are necessary for assembly but dispensable for silencing and peripheral positioning of heterochromatin

Reminiscent to the role of H1 in mammalian cells, Arabidopsis plant cells lacking the three canonical H1 isoforms show well constituted nuclei. Yet they exhibit a larger size and fail to form the typical 6-8 heterochromatic chromocenters (CC) normally seen in wild-type somatic nuclei (**Figure 1A,B**). Arabidopsis CCs are largely composed of centromeric and pericentromeric transposable element (TE) repeats and a subset of two to four CC are associated with the nucleolus and comprise rDNA repeats (Fransz, De Jong et al. 2002, Soppe, Jasencakova et al. 2002). While centromeric repeats are dispersed in *3h1* mutant nuclei, rDNA repeats localize in compact CC as in wild-type (**Figure 1C**), indicating that H1s are essential for maintaining structural, compact domains at (peri-) centromere regions but dispensable for the heterochromatinization of rDNA repeat loci. Interestingly, although lacking a canonical organization, centromeric repeats in *3h1* nuclei remain located at the periphery as described in wild type (Andrey, Kieu et al. 2010) (**Figure 1D**) suggesting that H1-mediated CC compaction occurs downstream of the spatial positioning of centromeric regions. High-resolution imaging further confirmed the presence in *3h1* nuclei of nanoscopic bodies of condensed chromatin possibly corresponding to dispersed heterochromatin regions that were not assembled into larger (chromocenter) structures (**Figure 1E**). The observation that H1s are required for CC formation is consistent with previous work showing that the Arabidopsis H1.1 variant is sufficient to induce ectopic heterochromatinization of genomic regions in tobacco (Prymakowska-Bosak, Przewloka et al. 1996). With a much shorter C-terminal tail, H1.3 shows a different chromatin-binding abilities, and a tissue-specific expression pattern distinct to that of H1.1 and H1.2 variants (Rutowicz, Puzio et al. 2015). Yet, the three Arabidopsis H1 variants can play a partially redundant function in CC organization in adult tissue. This is suggested by an intermediate reduction in heterochromatin content in the double *h1.1h1.2* mutant compared to that in the triple mutant in roots (**Figure 1 – figure supplement 1**). Possibly, the ectopic expression of H1.3 in the absence of H1.1 and H1.2 may contribute to this intermediate phenotype (**Figure 1 – figure supplement 1**). In embryonic tissues (cotyledons), CC formation and heterochromatinisation of centromeric and pericentromeric repeats is, however, clearly controlled by the H1.1 and H1.2 variants (**Figure 1 – figure supplement 1C**). Despite this functional redundancy, expression of an RFP-tagged H1.1 or GFP-tagged H1.2 variant is sufficient to restore heterochromatin assembly (**Figure 1 – figure supplement 1A,B**). At the genomic level, chromocenters display distinctive chromatin signatures described as chromatin states 8 and 9 (CS 8 and 9) and specifically enriched in H3.1 variants, DNA methylation, H3K27me1 and H3K9me2 modifications (Sequeira-Mendes, Araguez et al. 2014, Vergara and Gutierrez 2017). We generated chromatin accessibility analyses based on Micrococcal Nuclease profiling (MNase-seq) showing that typical CS8 and CS9 regions have a consistent 12-15% reduction in nucleosomal density in *3h1* nuclei (**Figure 1F**). In addition, nucleosome distribution is more variable in *3h1* heterochromatin as shown by the higher frequency of both short (<150nt) and unusually long (>300nt) MNase-protected regions compared to wild-type, with an average nucleosome repeat length NRL globally shorter by 10 nt in the *3h1* mutant (**Figure 1G**, **Figure 1 – figure supplement 2**). Thus, H1 seems to constrain nucleosomal spacing and provide a template of regularity in nucleosome distribution along heterochromatin regions, a property of H1 that was recently shown in Drosophila chromatin (Baldi, Krebs et al. 2018). This, in turn, may facilitate spatial folding into larger structures (Routh, Sandin et al. 2008) and hence chromocenter assembly. Furthermore, the absence of microscopically visible chromocenters does not seem to impair the deposition of their corresponding epigenetic silencing marks which remain abundant but massively redistributed in nucleoplasm (**Figure 1H**). Aiming at testing the effect of H1 on heterochromatin control, RNA-seq data were generated for both wild-type and *3h1* plants. Consistently, the transcriptional control of TEs still remains effective with only a moderate fraction of 1.5% TEs being upregulated in our RNAseq profiles (a third of them being LTR/Gypsy pericentromeric elements, **Tables S1 and S2, Figure 1 – figure supplement 3**). Collectively, our observations indicate that H1 is critical for heterochromatin assembly into compact chromocenter foci downstream of epigenetic silencing and peripheral positioning. This may be enabled through regulating nucleosomal distribution and constraining NRL ranges. Those define the density of nucleosomal arrays that in turn influence the folding and resulting structures into higher-order level chromatin arrangements (Maeshima, Rogge et al. 2016).

**Figure 1.**
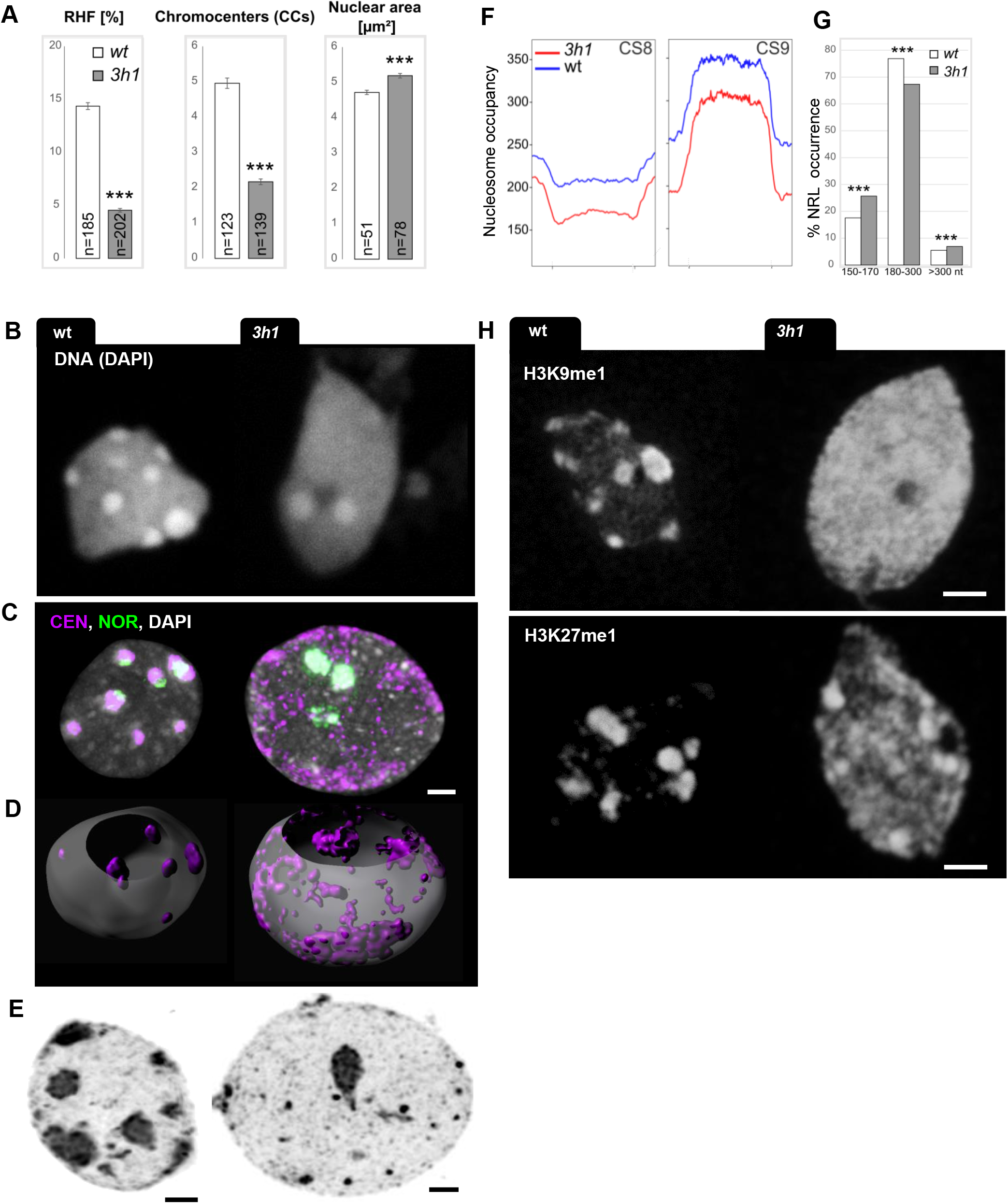
Loss of H1 variants leads to global chromatin decondensation but is dispensable for heterochromatin identity. Cytogenetic (**A-E, H**) and nucleosome profile (**F, G**) analyses of chromatin organization in triple *h1.1h1.2h1.3* (*3h1*) mutant and wild-type segregant (wt) seedlings. (**A**) H1 depletion induces a significant reduction of the relative heterochromatin fraction (RHF), the number of chromocenters (CCs) and an increase in nuclear size (area). ***, t-test, p-value < 0.001; error bars, standard error of the mean (s.e.m). Cytological analyses on isolated, spread leaf nuclei. (**B**) Typical wt and *3h1* nuclei as used in (A), stained with DAPI. (**C**) H1 depletion induces a spatial dispersion of the centromeric repeats (CEN, purple) but not the 45S rDNA, Nucleolar Organization Region repeats (NOR, green) as shown by Fluorescent *In Situ* Hybridization (FISH). (**D**) 3D segmentation of the CEN signals shows that the preferentially peripheral localisation of CEN repeats is unaffected in *3h1* nuclei despite their lack of condensation. (**E**) High-resolution imaging using STED microscopy and deconvolution-based reconstruction of *3h1* and wt nuclei. Nanoscopic bodies of condensed chromatin are dispersed throughout the nucleus in *3h1* instead of conspicuous chromocenters as in wt. (**F**) Nucleosome occupancy is lower in *3h1* heterochromatin, along the corresponding chromatin states 8 and 9 (Sequeira-Mendes, Araguez et al. 2014). (**G**) Distribution of Nucleosomal Repeat Lengths (NRLs) in wt and *3h1*, chi-square test, p-value < 0.0001 (***). (**H**) The heterochromatic marks H3K9me1 and H3K27me1 are not reduced but redistributed in *3h1* nuclei. Scale bar: 2μm. Isolated leaf nuclei were flow-sorted according to their 2C DNA content (A, B, E, H). The following figure supplements are available for figure 1: **Figure 1 – figure supplement 1**. Chromocenter formation relies on H1.1 and H1.2 but not H1.3 in differentiated, adult tissues. **Figure 1 – figure supplement 2.** H1 regulates chromatin-state dependent nucleosomal distribution. **Figure 1 – figure supplement 3.** Super families of TEs upregulated in *3h1* mutant.

### H1 variants enable a regular spatial distribution of nanoscale-chromatin domains and regulate nucleosomal density and mobility in euchromatin

As shown by genome-wide profiling, Arabidopsis H1 variants are abundant throughout the genome and, besides heterochromatin, are present in euchromatin regions (Rutowicz, Puzio et al. 2015). This is also visualized *in situ* (**Figure 2A**) showing discrete regions with enriched H1 levels interspersed with H2B (**Figure 2B**, **inset**). We thus hypothesized that H1 depletion may also impact the structural organization of euchromatin regions. To resolve nanoscale level–of-organization, we measured chromatin density patterns on ultrathin transmission electron microscopy (TEM) preparations (**Figure 2C**). For this we used a spatial pattern analysis approach that was previously validated to capture relevant, functional features of chromatin organization in cancerogenous animal cells (Cherkezyan, Stypula-Cyrus et al. 2014). In brief, a spatial autocorrelation function (ACF) of chromatin staining spatial distribution is calculated inside multiple regions of interests (ROIs, **Figure 2D, Figure 2 – figure supplement 1A**) within the euchromatin region of each nucleus, and is used to infer the distribution of structured signal intensities at given length scales (**Figure 2E**). Strikingly, the study unveiled that euchromatin of *3h1* nuclei harbors significantly less spatial homogeneity in nanodomain distribution, as shown by a less shallow autocorrelation fit in *3h1* (ACF, **Figure 2E**) and higher dispersion of length scales (*D*) compared to wildtype (**Figure 2F**). This trend was reversed in mutants complemented by a tagged H1.1 variant (**Figure 2 – figure supplement 1B**) and independently confirmed on super resolution microscopy images of fluorescently immunolabelled nucleosomes (**Figure 2 – figure supplement 1C**). Thus, H1 variants mediate both the organization and regularity of discrete, spatial nanodomains. At the molecular level, H1 depletion does not affect the overall qualitative distribution of nucleosomes with respect to chromatin states or metagene profiles in our MNase-seq analysis of *3h1* plants (**Figure 1 – figure supplement 2**). However, H1 depletion affects nucleosomal density, though in a variable manner, with regions showing higher coverage while others show no change or lower coverage. This is particularly well illustrated by nucleosome density profiles among chromatin states where the average levels relative to the CS boundaries are enhanced or, in contrast, diminished (for instance CS1, CS5 and CS4 states, **Figure 2G, Figure 1 – figure supplement 2**). This suggests that H1s provide structural attributes to epigenetically distinct domains. Next, we assessed whether the relaxation of chromatin domains in *3h1* influenced global nucleosomal mobility in euchromatin, a property that is strongly correlated with transcriptional competence in plants and animals (Schwabish and Struhl 2004). Fluorescence Recovery After Photobleaching (FRAP) was performed on cells expressing an RFP-tagged H2B variant showed that nucleosomes are ~2.5 times more mobile in H1-depleted chromatin than in wild-type (**Figure 2H**). Chromatin mobility in mutant differentiated cells resembled that in wild-type meristematic (pluripotent) cells (**Figure 2 – figure supplement 2**). Consistent with this higher mobility, *3h1* nuclei show global histone hyperacetylation typical for meristematic chromatin (Rosa, Ntoukakis et al. 2014), with a 2.5-fold increase at the cytological level compared to wild-type (**Figure 2I**).

**Figure 2.**
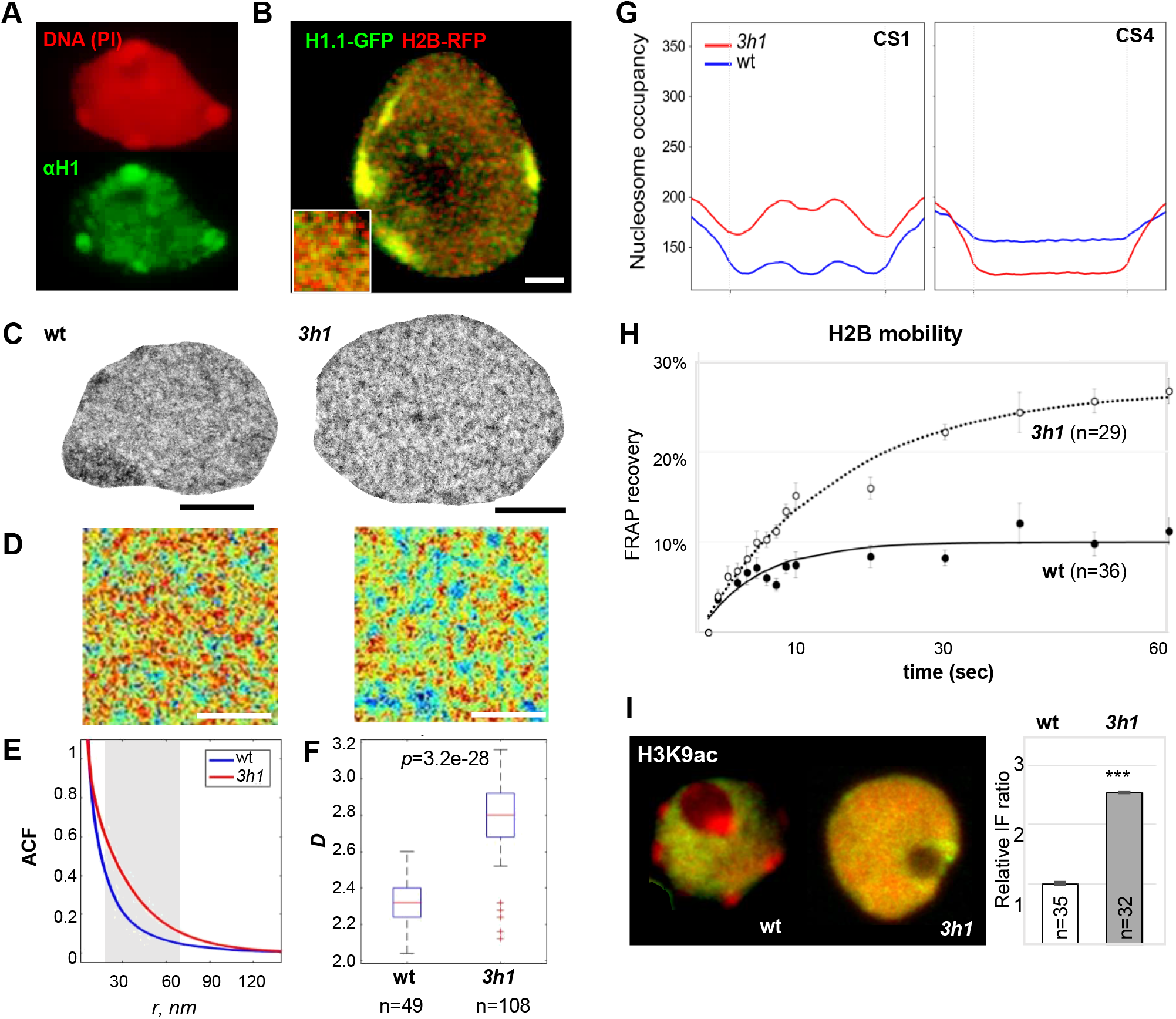
H1-depletion has a strong impact on euchromatin organization with increased dispersion of nanoscopic domains, altered distribution of nucleosome coverage and increased mobility. (**A-B**) H1 is abundant in euchromatin distributed as discrete foci partially colocalizing with H2B. (A) H1 immunostaining and Propidium Iodide (PI) counterstaining, done as in Figure 1, (B) live histone reporter imaging as indicated above the pictures. (**C-F**) Ultrastructural analysis of euchromatin organization in wt *vs 3h1*. (C) Typical TEM image of nuclei stained with uranylacetate on 7 nm cryosection (root epidermis, see Methods), Scale Bar: 1μm, (D) representative region of interest (ROI) in euchromatin of wt and *3h1* nuclei used for spatial autocorrelation function (ACF) analyses. Scale Bar: 500nm (E-F) Spatial chromatin density analyses show decreased regularity in the spatial chromatin distribution pattern in *3h1* revealed by a less shallow ACF curve within length scales of 20-60nm (grey zone, graph, E) and higher dispersion of length scales as shown by bigger range of the estimate *D* characterizing the spatial autocorrelation fit (F). These differences in *3h1* are restored upon complementation with an H1.1 expressing construct. ***, unpaired t-test, p-value < 0.001. See also **Figure 2 – figure supplement 1**. (**G**) Nucleosome coverage but not qualitative distribution is altered in H1 depleted euchromatin. Antagonist effects are seen for regions of chromatin states CS 1, 3, 7 (CS1 only is shown here) and CS4 (CS according to (Sequeira-Mendes, Araguez et al. 2014)). See also **Figure 1 – figure supplement 2** for nucleosome occupancy in *3h1* and wt over regions from all chromatin states. (**H-I**) Chromatin mobility is dramatically increased in *3h1* concomitantly to higher histone acetylation levels. (H), H2B-RFP fluorescence recovery in FRAP experiments, double normalisation, see Methods, (I) histone hyperacetylation in *3h1* leaf nuclei (experimental approach as in Figure 1). The following figure supplements are available for figure 2: **Figure 2 – figure supplement 1.** H1-depletion induces spatial dispersion of structural chromatin domains at the nanoscale level. **Figure 2 – figure supplement 2.** Chromatin mobility in meristematic nuclei is not affected by H1 depletion.

In conclusion, H1-depleted cells show a relaxed, hyperacetylated, highly mobile chromatin with a low degree of structural differentiation between chromatin states. Our data show that H1 variants play a significant role in euchromatin too where they modulate nucleosomal density, restrict nucleosome mobility and enable regularity at the spatial level in the distribution of higher-order, nanoscopic domains in the nucleus.

### H1 depletion alters epigenetic and structural signatures linked with transcriptional competence but *in fine* impacts only moderately gene expression

Next, we asked whether this global chromatin relaxation induced by H1 depletion would impact on the transcriptional landscape. In wild-type tissue, the nucleosomal coverage at genic regions inversely correlates with expression levels (**Figure 3A** and (Li, Liu et al. 2014)). Nucleosomal density in gene bodies, particularly (ie downstream the transcriptional start site, TSS) corresponds to a structural attribute distinguishing gene loci according to their transcriptional states. In *3h1*, we observed a notable loss of structural differentiation among these states with a generally higher nucleosomal density specifically downstream the TSS (**Figure 3A**). At the same time, though, this higher nucleosomal occupancy did not seem to impair transcription for a majority of genes since very few loci are downregulated in *3h1* plant lines (43 genes for p-value < 0.05 and fold change > 2, **Figure 3B, Table S3, Figure 3 – figure supplement 1**). By contrast, most of the moderate fraction of genes that are misexpressed in *3h1* are up-regulated (658 genes for p-value < 0.05 and fold change > 2, **Figure 3B, Table S3**). Therefore, H1 is necessary to provide distinct structural signatures to genomic regions with distinct transcriptional profiles, but does not affect transcriptional competence at a global level. Yet, H1s clearly exert a transcriptional control at a few hundreds of loci. Interestingly, down-regulated genes are largely representing light-related metabolism with an enrichment in Gene Ontology (GO) terms related to chlorophyll, photosynthesis and response to light (**Table S4**). We did not find a specific enrichment in GO terms for the group of up-regulated genes (not shown), nor a dramatic overrepresentation of specific chromatin states (**Figure 3 – figure supplement 3**). However, an interesting observation is that these genes have a notable high periodicity in nucleosome positioning within 800bp downstream the transcriptional start site (TSS, **Figure 3C**), a feature which is normally only found for a subset of (highly) expressed genes (Pass, Sornay et al. 2017), whereas this class of H1 targets are low expressed in wildtype (**Figure 3 – figure supplement 2**).

**Figure 3.**
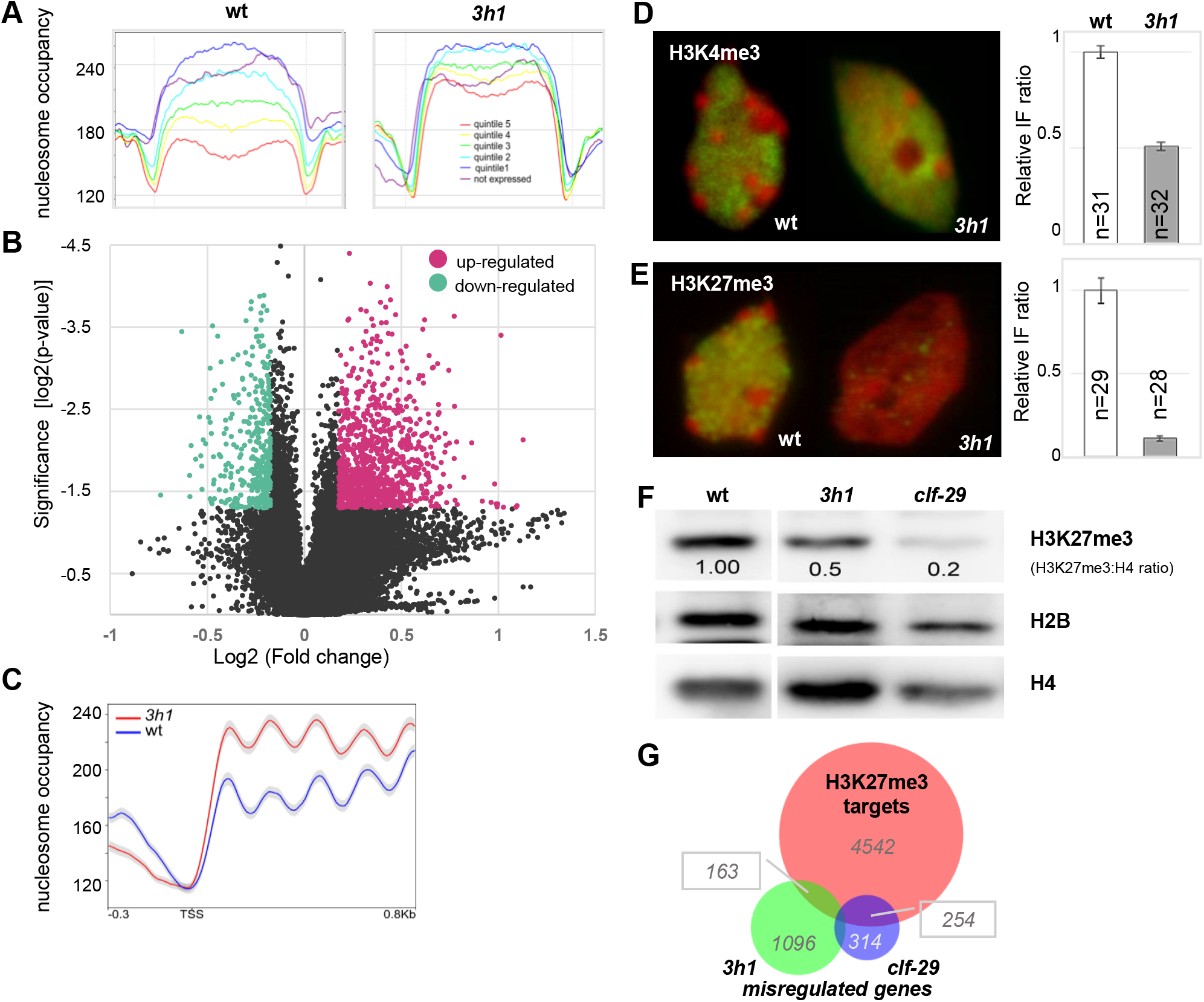
H1 is necessary to secure transcriptional state-specific nucleosomal and epigenetic profiles yet influence only a moderate gene fraction. (**A**) Nucleosome distribution profiles clearly defines distinct gene classes according to expression levels in wild-type but no longer in *3h1*. Quintiles 5 to 1 represent categories of genes with expression levels ranked from the highest to lowest level, respectively, as previously described (Rutowicz, Puzio et al. 2015). (**B**) H1 depletion induces moderate changes in the transcriptional profile yet a subset of 701 genes (p-value < 0.05 and fold change > 2) are misregulated. The volcano plot was cropped around the denser part of the dataset. The full plot is presented in Figure 2 – figure supplement 1. (**C**) Up regulated loci show a characteristic nnucleosome occupancy with high periodicity and a higher coverage in *3h1* downstream the TSS. TSS, Transcription Start Site. (**D-E**) Decreased abundance of H3K27 and H3K4 trimethylation in *3h1* measured by quantitative immunostaining on isolated leaf nuclei. (**F**) A two-fold reduction of H3K27me3 levels upon H1 depletion is confirmed by Western-Blot on seedling leaves, yet is less dramatic than in a loss of PCR2 function mutant, *clf-29*.The original picture is presented in Figure 2 – figure supplement 2. (**G**) Genes which are up-regulated in *3h1* share a significant overlap with H3K27me3 targets defined by (Zhang, Clarenz et al. 2007) (P=0.0007, Fisher exact test) but remain distinct from those affected by the *clf-29* mutation (Wang, Liu et al. 2016). Legend: (**D**), (**E**) green, immunostaining; red, DNA counterstaining; graph, relative fluorescence intensity – antibody signals normalised over DNA content (see Methods). The following figure supplements are available for figure 3: **Figure 3 – figure supplement 1.** Gene expression in *3h1* mutant. **Figure 3 – figure supplement 2.** Up‐ and down-regulated genes in *3h1* correspond to gene categories with distinct expression strength in wild-type. **Figure 3 – figure supplement 3.** Distribution of chromatin states (CS) of all genes in the *Arabidopsis thaliana* genome versus genes upregulated in *3h1* plants. **Figure 3 – figure supplement 4.** The overall distribution of CHH and CG is not affected in *3h1* mutant nuclei. **Figure 3 – figure supplement 5.** H3, H3K4me2 and H3K27me2 are not affected by H1 depletion. **Figure 3 – figure supplement 6.** Global levels of H3K27me3 are reduced in *3h1* seedlings.

The impact of H1 depletion on gene expression may arise from improper nucleosome distribution in regulatory regions influencing the access of the transcription machinery and epigenetic regulators as this was shown for DNA methylation (Wierzbicki and Jerzmanowski 2005, Zemach, Kim et al. 2013, Lyons and Zilberman 2017). In addition to influence on DNA methylation, H1 depletion was also shown to correlate with drastic changes of the histone modification landscape in the context of germline precursor (Spore Mother Cells, SMC) differentiation: there, H1 eviction is a developmental marker of the somatic-to-reproductive fate transition that precedes a breadth of global chromatin changes at the structural and epigenetic levels (She, Grimanelli et al. 2013). These include heterochromatin decondensation and histone hyperacetylation as seen in *3h1* mutant somatic tissues as well as a marked elevation of H3K4me3 and decrease of H3K27me3 levels, respectively (She, Grimanelli et al. 2013), together with a transient decrease of DNA methylation in the CHH, but not the CG sequence context (Ingouff, Selles et al. 2017). We thus looked at the cytological distribution and abundance of DNA methylation and canonical chromatin modifications in *3h1* mutant nuclei (**Figure 3 and Figure 3 – figure supplements 4 and 5**). Cytological levels of methylated DNA in the CG and CHH context was not altered in *3h1* mutant nuclei (**Figure 3 – figure supplement 4**) which indicates that genome-wide erasure of CHH DNA methylation in SMC (Ingouff, Selles et al. 2017) is not simply a consequence of H1 depletion. In addition, H1 was shown to influence the DNA methylation landscape in a complex manner depending on other genomic and chromatin attributes (Wierzbicki and Jerzmanowski 2005, Zemach, Kim et al. 2013), which cannot be captured by cytological imaging. H3K4me3 are moderately, but reproducibly lower in *3h1* nuclei (**Figure 3D**). Thus, H3K4me maintenance in somatic tissues requires H1. A corollary to this is that chromatin decondensation is not the cause of H3K4 hypermethylation in H1-depleted SMC as initially interpreted. In addition, H1 depletion in *3h1* mutant nuclei resulted in a drastic reduction of H3K27me3 levels compared to wild-type (**Figure 3E**) that was originally measured in H1-depleted SMCs (She, Grimanelli et al. 2013), but not of H3K27me2 (**Figure 3 – figure supplement 5**). A two-fold reduction of H3K27me3 was further confirmed on whole seedling chromatin extracts by immunoblotting (**Figure 3F and Figure 3 – figure supplement 6**). Yet, only 10% of the genes upregulated in *3h1* (p-value<0.05) overlap with known H3K27me3 genomic targets, and ca 4% are shared with a PRC2 compromised mutant such as *clf* (*curlyleaf*, (Wang, Liu et al. 2016)). Thus, although these proportions remain significant when compared to random representations (p-value<0.001, Fisher test) it suggests that loss of H3K27me3 is not solely responsible for gene misregulation in *3h1* seedlings. The expression of histone modifying enzymes is not significantly changed in *3h1* (**Table S5**). This indicates that altered H3K4me3 and H3K27me3 landscapes may rather be a consequence of altered nucleosomal density, as shown before, possibly affecting the targeting, spreading of the modifications, or both.

Collectively, our data indicate that H1 variants provide structural attributes enabling differentiation of transcriptional domains and maintenance of histone modifications in euchromatin. Although these attributes are not essential with regards to plant growth under laboratory conditions, our analyses unveil that H1s are required for transcriptional control at several hundred loci.

### H1 reinforces the epigenetic controls of developmental and cellular transitions

*3h1* mutant plants resume a functional organism suggesting that H1-mediated chromatin organization is dispensable for the basic functioning of the plant genome in laboratory growth conditions. However, we observed several subtle but quantifiable deviations from the otherwise highly regular developmental pattern observed in the wild type. Particularly, *3h1* plants were affected at key developmental transitions of the plant’s life cycle such as seed dormancy breaking and flowering (**Figure 4A,B**) as well as during cellular transitions in root and shoot tissues responsible for the establishment of lateral roots, root hairs and leaf stomata (**Figure 4C-F**). More specifically, *3h1* seeds showed a prolonged seed dormancy (**Figure 4A**) where the expression of H1.1 was sufficient to restore a wild-type trait. Following germination, mutant plants grew regularly but flowered significantly earlier (**Figure 4B**), a rescuable phenotype mostly attributed to H1.1 and H1.2 variants (**Figure 4 – figure supplement 1**).

**Figure 4.**
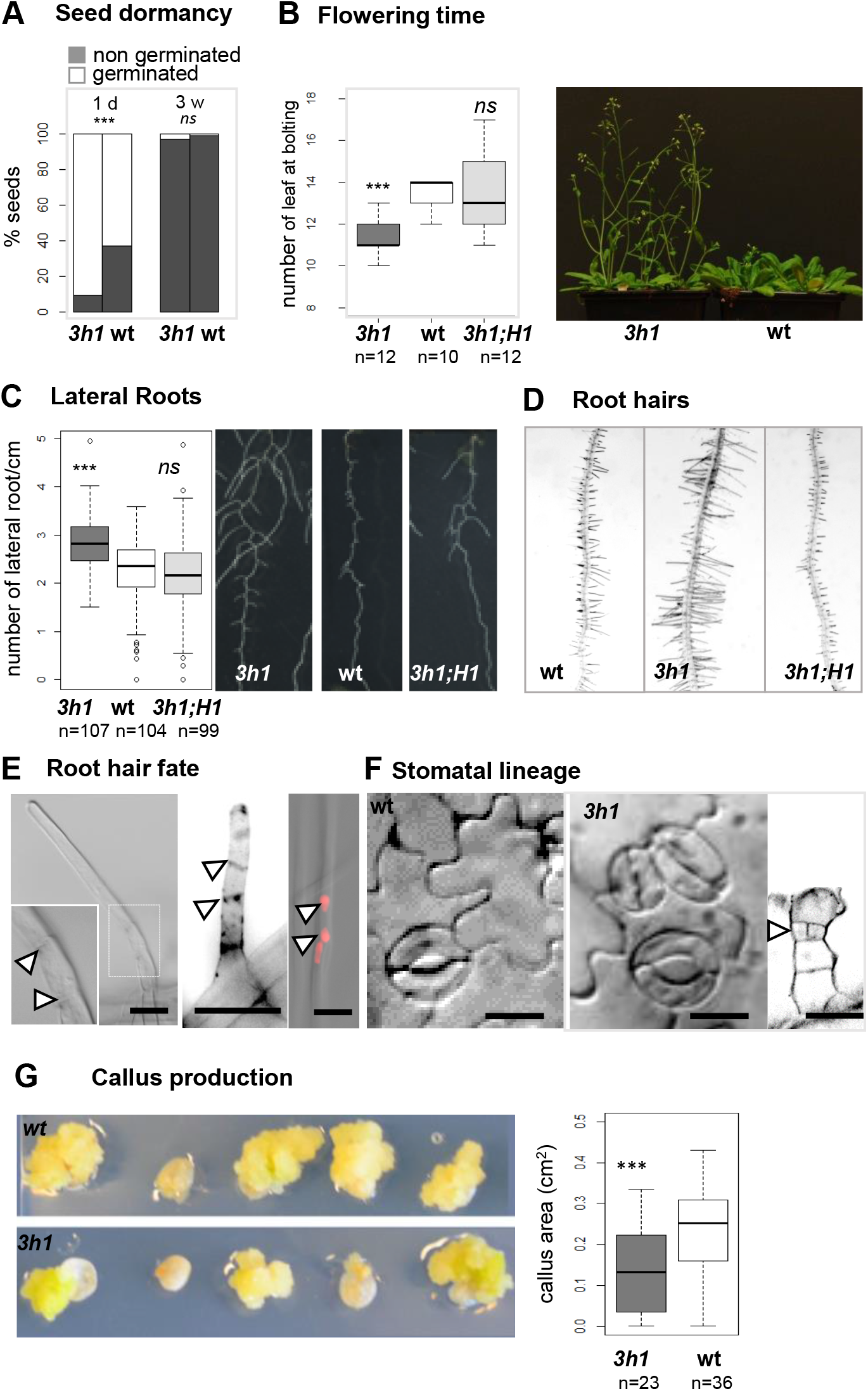
H1-depletion relaxes the epigenetic control of several phase and cellular transitions. The *3h1* mutant shows relaxed control of seed dormancy (**A**), flowering time (**B**), lateral root formation (**C**), root hair density (**D**) and fate (**E**), stomatal spacing (**F**) and is impaired reprogramming competence *in vitro. 3h1* shows, compared to wild-type (**A**) a delayed seed germination competence in mature seeds 1 day post harvest but not in dried seeds (3 weeks post harvest), (**B**) early flowering measured as the number of rosette leaves at bolting, (**C**) increased number of lateral roots (8 DAG seedlings), (**D**) increased root hair density, (**E**) occasional multicellular root hairs, (**F**) stomatal complexes with reduced spacing and supernumerary divisions of the lineage precursor (adaxial cotyledon epidermis, 10 DAG; DIC pictures for wt and *3h1* (*middle*), Renaissance counterstaining (*3h1′*, *right*)), and (**G**) decreased callus size under induction medium. Wild-type segregants (wt) were compared with triple mutant tissues/seedlings (*3h1*) and, whenever indicated, with complemented lines expressing H1.1 and H1.2 variants only (*3h1*; H1) or all three variants (*3h1*; H1*). Statistical tests (A,B Welch t.test; C, Fisher exact test) were performed against wt replicates, *** p-value < 0.001, ns, not significant. The following figure supplements are available for figure 4: **Figure 4 – figure supplement 1.** Flowering time for h1 mutants and complemented lines. **Figure 4 – figure supplement 2.** H1 is required for correct developmental transitions. This figure shows additional image and quantification material supporting Figure 4 **Figure 4 – figure supplement 3.** Callus formation efficiency in H1 deficient mutants is reduced. **Figure 4 – figure supplement 4.** H1.2 levels decrease relative to H1.1 during cellular differentiation in root.

Besides these major developmental transitions changing the plant’s lifestyle, several cellular transitions occur that establish tissues and cell types that are not predefined in the primary root or shoot organs, hence are not a meristem-derived lineage. In Arabidopsis, the specification of lateral root primordia from pericycle founder cells and the differentiation of root hairs from epidermal cells follow a regular pattern modulated by developmental and environmental cues (Van Norman, Xuan et al. 2013, Salazar-Henao, Velez-Bermudez et al. 2016). Compared to wild-type, *3h1* seedlings produced more lateral roots per root length unit (**Figure 4C, Figure 4 – figure supplement 2**), more root hair cells (**Figure 4D**, **Figure 4 – figure supplement 2**). Both phenotypes were reversed upon restored expression of H1.1 and H1.2, possibly indicating more frequent developmental initiation events. In addition, the unicellularity of root hairs was occasionally compromised in *3h1* with the appearance of multiple nuclei and cell boundaries (**Figure 4E, Figure 4 – figure supplement 2**). Similarly, stomata patterning was altered in *3h1* mutant leaf epidermis with a higher occurrence of high-degree (tertiary and quaternary) clusters, associated with complex arrangements, collated stomata or atypical division patterns in early stages that were not found in the wild type (**Figure 4F, Figure 4 – figure supplement 2**). These observations suggest a loose control of stomatal spacing presumably involving occasional re-initiation events (Lau and Bergmann 2012). Finally, we also tested how *3h1* tissues respond to reprogramming in *in vitro* culture and indeed measured a decreased efficiency in callus development compared to the wild-type (**Figure 4G**), a feature mostly attributed here to H1.3 (**Figure 4F, Figure 4 – figure supplement 3**). Interestingly, all these phenotypes point out to processes regulated by PRC2 complexes as demonstrated by genetic analyses or inferred from PRC2-mediated enrichment of H3K27me3 at regulatory loci controlling these transitions (Wood, Robertson et al. 2006, Bouyer, Roudier et al. 2011, He, Chen et al. 2012, Gu, Xu et al. 2014, Lee, Lucas et al. 2014, Molitor, Bu et al. 2014, Zhu, Rosa et al. 2015). Not all three H1 variants are equivalently involved in these processes, with H1.1 and H1.2 largely contributing to flowering, lateral roots and dormancy while H1.3 may be solely responsible for callus competence. Although relative levels of H1.1 and H1.2 variants change along the meristematic-elongation-differentiation transition in roots (with an increasing H1.2/H1.1 ratio, (**Figure 4F, Figure 4 – figure supplement 4**) the moderate but specific phenotypic alterations in *3h1* mutant plants suggest a relaxation of some of the mechanisms controlling cellular transitions, possibly as a consequence of the singular chromatin organization in this mutant.

## Discussion

H1 linker histones are core components of chromatin organization in eukaryotes. Thanks to a tripartite structure, H1s bind the DNA at entry/exit sites and tether neighboring octamers through electrostatic interactions with positively charged, flexible tails. As a result, H1s spatially accommodate a string of nucleosomal particles in an ordered, spatial arrangement of a compaction level varying depending on H1 subtypes. Formerly proposed to achieve the folding of chromatin fibers into a large-scale, 30nm diameter solenoid, H1s are now understood to foster the formation of local nucleosomal arrays (also called ‘nucleosome clutches’ or ‘nucleosome clusters’) of varying size and density along the genome and depending on cell type (Ricci, Manzo et al. 2015) as shown in animal models. Reduction of H1 levels in mammalian cell lines generate cytological and molecular alterations of chromatin organization with heterochromatin reorganization, nuclear swelling, loss of sharp boundaries between topological domains, reduction of the average nucleosomal length repeat and loss of periodicity in nucleosomal array organization, alteration of the level and distribution patterns in histone modifications and DNA methylation (Fan, Nikitina et al. 2005, Cao, Lailler et al. 2013, Geeven, Zhu et al. 2015, Baldi, Krebs et al. 2018, Fyodorov, Zhou et al. 2018). Flowering plants and animals diverged more than 1500 MY ago and H1 protein sequence have substantially evolved in the different kingdoms, while their tripartite constitution remained preserved (Kasinsky, Lewis et al. 2001, Jerzmanowski 2007, Kotlinski, Knizewski et al. 2017). Functional similarities between plant and animal H1’s was shown for a few features, including the property to induce heterochromatin formation *in vivo* (Prymakowska-Bosak, Przewloka et al. 1996) and influence the DNA methylation landscape (Wierzbicki and Jerzmanowski 2005, Rea, Zheng et al. 2012, Zemach, Kim et al. 2013). Yet so far, a detailed analysis of H1 roles on chromatin organization in plants was missing. Here, we show that Arabidopsis variants collectively contribute to the fine-scale (nucleosome distribution, nanoscopic domains) and nuclear-scale (microscopic domains, global properties) levels of chromatin organization, with notably distinctive functions in eu‐ and heterochromatin.

Our cytological and expression analysis shows that for heterochromatin, H1 seems to act downstream the processes securing epigenetic silencing at TEs, notably repressive DNA methylation and H3K9 methylation, and peripheral positioning. TE close to genes, and located mostly along chromosome arms, require the RNA-dependent DNA Methylation (RdDM) machinery involving small RNAs for initial DNA methylation targeting; silencing maintenance is then reinforced by H3K9 methylation and DNA methylation maintenance enzyme (reviewed in (Sigman and Slotkin 2016)). By contrast, DNA methylation of pericentromeric TEs is independent of RdDM but requires CMT2, a specific DNA methyltransferase and the chromatin remodeler DECREASE in DNA METHYLATION1, DDM1 (Zemach, Kim et al. 2013). At pericentromeric TE loci, H1 is thought to modulate (but not hinder) DNA methylation in the CHG context by reducing access to CMT2, a configuration resolved by DDM1 (Zemach, Kim et al. 2013). Interestingly, despite a globally stable TE expression landscape in *3h1*, a small fraction (1.5%) of TEs, which are mostly pericentromeric (enriched in LINE, Gypsy and Copia elements) is derepressed upon H1 depletion. Thus, this subset of TE might reveal a dual role for H1 in possibly a cooperative role here (instead of hindrance) with CMT2 for establishing a repressive DNA methylation profile. Alternatively for this subset of TEs, H1 act in a distinct manner together with CMT3 with which it was shown to interacts (Du, Zhong et al. 2012). In mammalian stem cells, H1 depletion leads to the derepression of major, pericentromeric satellite repeats (but not of the centric, minor satellite repeats) independently of common epigenetic silencing marks (Cao, Lailler et al. 2013). In Drosophila, however, H1 depletion also releases silencing of pericentromeric TEs but in an H3K9me2-dependant manner (Lu, Wontakal et al. 2013, Iwasaki, Murano et al. 2016). Thus, while the situation in Arabidopsis seems closer to that described in mammalian cells, Arabidopsis H1 may share a dual role in in modulating either negatively or positively pericentromeric repeats silencing.

At the nuclear level, we show here that H1 variants are necessary to assemble marked, silent and positioned heterochromatic regions into compact, microscopic structures known as chromocenters (Figure 5). On the other hand, in Drosophila, particularly in salivary gland cells H1 is responsible for pericentromeric heterochromatin assembly (Lu, Wontakal et al. 2009), but this might be considered as a singular situation due to polytenic chromosomes. It thus remains that animal and plant H1 subtypes may differ in their function regarding chromocenter assembly and spatial arrangement in interphase nuclei. In addition and interestingly, genomic repeats of the NOR are not affected by the loss of H1 variants in Arabidopsis nuclei indicating the existence of an H1-independent control of these heterochromatic structures. Which factors, possibly among GH1-containing proteins (Kotlinski, Knizewski et al. 2017), mediate the compaction of NOR in H1 depleted cells remains to be determined.

**Figure 5.**
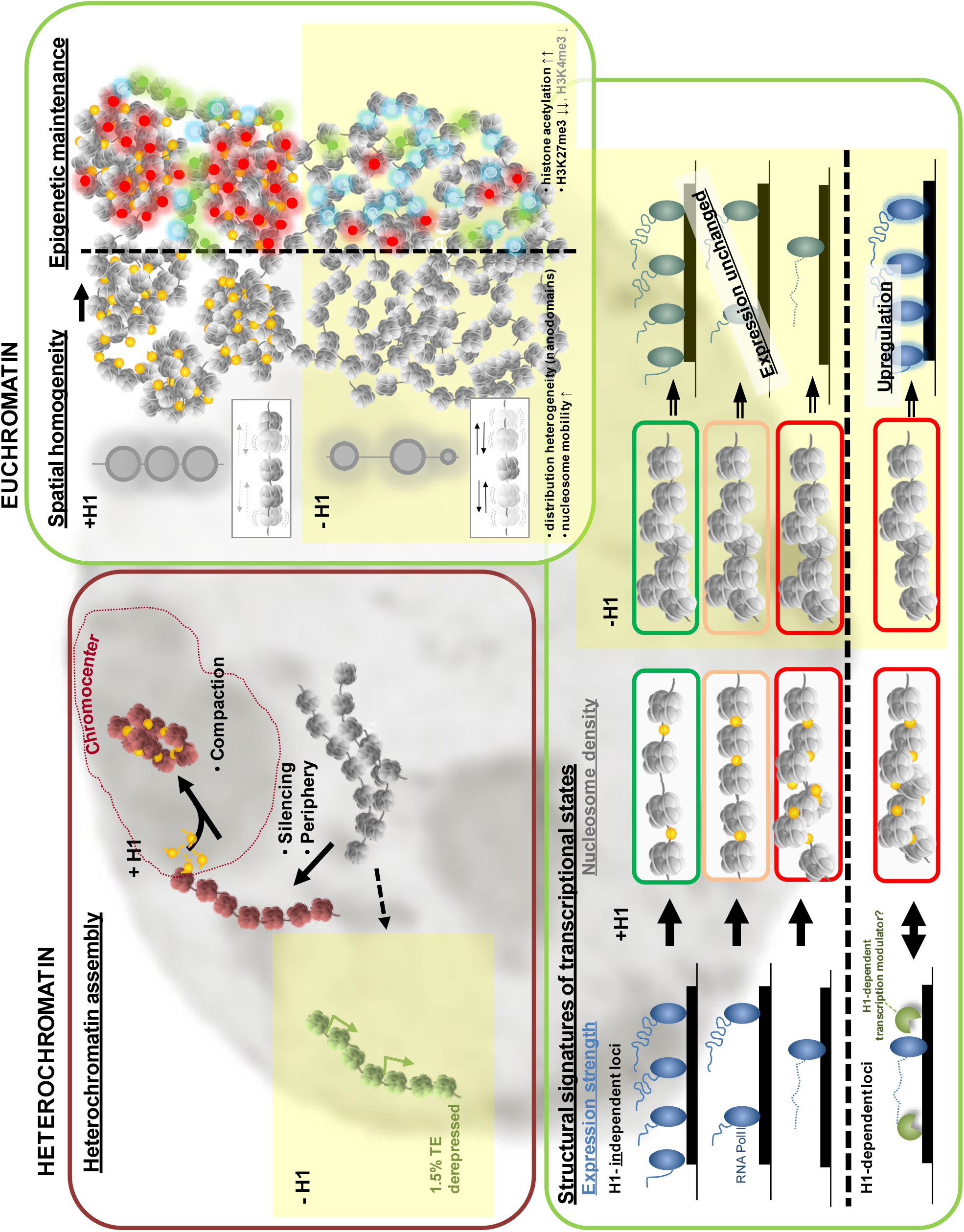
Model for H1 function in heterochromatin and euchromatin organization at the topological and molecular level. Graphical representation of H1 roles on chromatin organization at the cytological (spatial) and molecular level based on analyses reported in this study. <u>Heterochromatin</u>: H1 is dispensable for silencing and peripheral positioning of the vast majority of heterochromatic repeats but necessary for their molecular assembly into compact chromocenter domains; yet a subset of Transposable Elements are directly affected by H1 and become derepressed in its absence (yellow box, -H1). This indicates both H1-independant and H1-dependent TE silencing controls. <u>Euchromatin</u>: Top right panel, H1 is necessary to provide homogeneity in chromatin topology and spatial organization of chromatin domains. H1 depletion results in both larger gaps between nanodomains, possibly enabling increased accessibility, and irregular, high local compaction; this chromatin heterogeneity is reminiscent of H1-depleted pluripotent cells (Ricci et al, 2015), cells with a loss of a SWI/SNF chromatin remodel function or undergoing tumorigenic reprogramming (Almassalha, Tiwari et al. 2017). Concomitantly, H1 depleted chromatin displays histone hyperacetylation (blue), increased mobility and poor maintenance of histone H3 lysine 4 (green) and more strongly lysine 27 (red) methylation. At the molecular level (lower panel), H1 provides distinct structural signatures (nucleosome coverage) at loci marked by distinct expression rates but is not epistatic to transcriptional control for a majority of them (H1-independant regulation); a subset of genes (ca 600 under a stringent cut-off), however, display an H1-dependent control possibly involving transcriptional regulators directly influenced by H1.

In euchromatin, the spatial distribution of chromatin density patterns becomes heterogeneous in H1-depleted chromatin compared to wild-type nuclei. Autocorrelation analyses showed that *3h1* chromatin is a more heterogeneous material composed of irregularly dispersed, nanoscale patches of variable densities as opposed to wild-type. This spatial dispersion is accompanied by an altered distribution of key histone 3 methylation marks which raises a question of causality between these two phenotypes (Figure 5). Nucleosomal arrays can be formed in the absence of H1 yet with less regularity, forming “ladders” or “puddles” type of arrangements (Beshnova, Cherstvy et al. 2014). The increased spatial dispersion of compact nanoscopic chromatin domains as observed in TEM in *3h1* nuclei is highly reminiscent of euchromatin distribution in tumorigenic nuclei losing fractal property of organization (Cherkezyan, Stypula-Cyrus et al. 2014). These specific disturbance in spatial chromatin organization have been correlated with transcriptional heterogeneity which are explained by the paired effect of increased accessibility and increased local compaction (Almassalha, Tiwari et al. 2017) similarly to the situation in *3h1*. Our observations are also consistent with reports that lower levels of H1 in pluripotent mammalian cells are responsible for changes in spatial chromatin distribution patterns with higher dispersion of nucleosome clutches with of smaller size, increasing PolII accessibility and favoring its redistribution (Ricci, Manzo et al. 2015).

At the molecular level, H1 influences both nucleosomal spacing and abundance with, however, distinct and sometimes antagonistic consequences depending on chromatin states. Notably, nucleosomal coverage is smaller in H1-depleted chromatin relative to wild-type in states typical for no or poor transcriptional activity (intergenic regions, heterochromatin regions). Yet this change in nucleosomal distribution does not correlate with a global derepression of the corresponding genic regions reinforcing the idea that H1 acts downstream of processes maintaining a durable epigenetic silencing (as opposed to other, variably expressed loci) in Arabidopsis. By contrast to transcriptionally silent regions, H1 depletion induced an increased level of nucleosomes at chromatin states/cis genomic elements normally associated with transcriptional competence without compromising transcription for a vast majority of loci (Figure 5). However, a few hundred loci are more directly influenced by H1 and linker histone depletion result in their upregulation concomitantly to a 1.2-fold nucleosome enrichment in their gene body-but not upstream regions-(Figure 5). These observations challenge an intuitive expectation coming from reports that nucleosome occupancy is usually inversely correlated to gene expression level (Li, Liu et al. 2014).

It indicates instead, that, at least in the absence of H1, a higher density of nucleosomes does not hinder transcriptional processes *per se*. Increased nucleosome mobility associated with higher histone acetylation in the absence of H1 might provide a functional chromatin template for transcriptional processes to be operated normally throughout the genome, except at a few hundred loci that seem H1-dependent. Upregulation in *3h1* is notably affecting genes that are normally very low or not expressed. Thus, this observation suggests two classes of repressed genes, those where H1-mediated chromatin organization is epistatic to transcriptional repression and those where H1 acts downstream the regulatory processes. This proposed explanation should be, however, pondered by the possibility that upregulation occurs only in a few cells, which chromatin profile is not captured in bulk tissue profiling. Nevertheless, our observations are reminiscent of the moderate, but not null, impact of H1 depletion in gene expression in animal cells (Shen and Gorovsky 1996, Hellauer, Sirard et al. 2001, Fan, Nikitina et al. 2005, Sancho, Diani et al. 2008) suggesting as well the existence of both H1-dependent and independent transcriptional control (**Figure 5**).

We further show that H1 depletion in Arabidopsis alters nucleosomal spacing but in a distinct manner from that in animal cells: besides a shift of NRL distribution peak from 170-180 to 160-170bp which is similar to the situation in animals (Fan, Nikitina et al. 2005), *3h1* chromatin shows a higher representation of both long (>200bp) and short NRLs (<160bp). This indicates a more permissive environment enabling diverse configuration of nucleosome spacing normally rare in the presence of H1 (Beshnova, Cherstvy et al. 2014). This possibly reflects the sterical influence of H1 imposing a defined range of nucleosome clustering, in turn influencing chromatin fiber folding and compaction (Routh, Sandin et al. 2008, Correll, Schubert et al. 2012). NRL variation is recognized to contribute chromatin fiber polymorphisms (reviewed in (Boule, Mozziconacci et al. 2015)). Long NRL can be composed of closely associated nucleosomes, but also of “stretched nucleosomes” formed in the absence of H1 and where DNA-histone contacts are lose (Usachenko, Gavin et al. 1996)). These molecular scenarios do not exclude the possibility that alternative chromatin architects such as HMG proteins (Jerzmanowski, Przewtoka et al. 2000, Postnikov and Bustin 2016) or H1-related proteins (Kotlinski, Knizewski et al. 2017) contribute to regulate nucleosome distribution and shape chromatin domains in *3h1* nuclei.

The consequences of H1 depletion at microscopic and ultrastructural level of chromatin organization in Arabidopsis is reminiscent of the pluripotent chromatin state in mammalian cells (Boskovic, Eid et al. 2014, Ricci, Manzo et al. 2015) but also of that of the Arabidopsis SMCs which are functional equivalent of the animal primordial germ cells. Indeed, H1 variants in Arabidopsis are actively depleted in both male and female SMC that undergo the somatic-to-reproductive cell fate transition. H1 eviction occurs at the onset of the meiotic S-phase and precedes nuclear enlargement, reduction in heterochromatin content (without affecting the level of the typical TE silencing H3K9me2 mark) along with drastic changes in the level and distribution of histone modifications in euchromatin with respect to canonical permissive and repressive histone marks (She, Grimanelli et al. 2013, She and Baroux 2015). Notably, H1 depletion in SMC was followed by an increase in H3K4 methylation and H4 acetylation. We observed here a corresponding hyperacetylation in *3h1* mutant cells similar to the one reported in SMC. This similarity is consistent with the known function of H1 in repressing core histone acetylation, likely by masking the target histone tail residues (Herrera, West et al. 2000). By contrast, opposite to what we observed in H1-depleted SMC, global levels of H3K4me3 are decreased in *3h1* mutant cells. Mammalian cells with reduced H1 levels showed a global increase of H3K4 methylation (Wang, Paucek et al. 2017) although a few loci loci showed lower levels (Geeven, Zhu et al. 2015). In H1 depleted SMC cells, increased H3K4 methylation is requiring the SET-domain H3K4 methyltransferase SDG2 (She, Grimanelli et al. 2013). But here, expression of most HMT-encoding genes are unaffected in the *3h1* mutant suggesting that H3K4me3 depletion is a primary effect of altered chromatin organization. Furthermore, developmentally-regulated depletion of H1 in SMC also had a considerable effect on H3K27me3 levels, similar to the drastic reduction we observe in the *3h1* mutant and to what was formerly reported in H1-depleted animal cells on pluripotency genes (Zhang, Cooke et al. 2012). Thus, most likely the dependence of the PRC2 complex towards H1-containing oligonucleosomes for propagating H3K27 methylation described in mammalian cells is conserved in plants (Martin, Cao et al. 2006). Despite a significant enrichment of *3h1*–misregulated genes in H3K27me3 targets, expression of only a fraction of the PRC2-regulated loci (profiled in (Wang, Liu et al. 2016)) are affected by H1 depletion indicating that at least in Arabidopsis, there are alternatives to H1-mediated chromatin structures for PRC2 activity. Conversely, ectopic transcription, enabled at H1-depleted loci, may also provide a mechanism for PRC2 inhibition as suggested previously (Wang, Paucek et al. 2017).

Collectively, our data show that chromatin dynamics in plant and animal cells share common organizing principles in fine-scale and nuclear-scale level of chromatin organization orchestrated by linker histone variants, and in the influence of H1 on the epigenetic landscape notably in euchromatin. Given an early origin of H1-related histones and PRC2 components (Kasinsky, Lewis et al. 2001, Shaver, Casas-Mollano et al. 2010), the functional relationships between H1-mediated chromatin structure and PRC2 activity possibly predate the dichotomy between plant and animal kingdoms. The evolutionary conservation of H1 function as architect and epigenetic modulator is however in stark contrast with their apparent dispensability, as many organisms tolerate H1 depletion (Izzo and Schneider 2016). H1 depletion has a moderate impact on gene expression at a global scale in Tetrahymena, yeast, vertebrates (reviewed in (Izzo and Schneider 2016)) and plants (this study) suggesting that H1 is not epistatic over basic molecular controls of transcription, at least for a large fraction of the genome. Nonetheless, in both plant (this study) and animal (Lu, Wontakal et al. 2013, Geeven, Zhu et al. 2015) cells, several hundred loci are misregulated upon H1 depletion including a large fraction of upregulated genes which transcriptional repression is thus normally H1-dependent.

The observation that H1 depletion in mammalian stem cells affect pluripotency genes (Zhang, Cooke et al. 2012), that H1 variants are evicted in a developmentally-regulated manner in primordial germ cells prior to pluripotency establishment in mouse (Hajkova, Ancelin et al. 2008) but also in plants in the precursors of the male and female reproductive lineage (She, Grimanelli et al. 2013, She and Baroux 2015), and that H1 incorporation is reduced in pluripotent mammalian cells (Ricci et al 2015) suggest a role for H1 in establishing a chromatin environment favorable to epigenetic and transcriptional reprogramming. The subtle yet specific phenotypes of *3h1* in the flowering and dormancy transitions, in the control of lateral root formation, root hair and stomatal fates, in *in vitro*-induced tissue reprogramming (callus) fuel the hypothesis that H1-mediated chromatin organization may facilitate epigenetic reorientation during cellular transitions. Consistent with this concept, H1-depleted nuclei show chromatin organization and properties to some extent reminiscent of an “immature” state of chromatin organization typical of pluripotent plant cells and/or meristematic tissues (She, Grimanelli et al. 2013) (Tessadori et al, 2010, Costa et al 2014) and pluripotent cells in animals (Boskovic, Eid et al. 2014, Ricci, Manzo et al. 2015). These consideration prompts for further investigations aiming at testing the specific role of H1 in gene expression robustness *versus* variability (Cortijo, Aydin et al. 2018), especially during environmental challenge frequent in natural condition. Furthermore, the connection between spatial dispersion in chromatin patterns distribution and transcriptional heterogeneities in H1-depleted plant cells echoes with recent models in mammalian cells implicating higher-order folding chromatin topology as an independent route influencing transcriptional dynamics (Almassalha, Tiwari et al. 2017). These evidence strongly motivate further exploration of fine-scale, spatial chromatin dynamics complementary to molecular-level, epigenetic studies of plant developmental and environmental response processes.

## Materials and Methods

### Plant materials and growth conditions

The *Arabidopsis thaliana* plants used in all experiments were in the Col-0 background unless it is specified otherwise. The *h1.1h1.2h1.3* (*3h1*) mutant was described before (She, Grimanelli et al. 2013) and it showed no detectable levels of H1 in Western Blot and immunostaining experiments (She, Grimanelli et al. 2013), (www.agrisera.com/en/artiklar/h1-histone-h1.html). Complemented mutant lines were generated by transforming *3h1* via floral dip method (Clough and Bent 1998) with H1 tagged variants (prom.H1.1::H1.1-RFP, prom.H1.2::H1.1-(G/C)FP, prom.H1.3::H1.3-GFP) described previously (She, Grimanelli et al. 2013, Rutowicz, Puzio et al. 2015). The *3h1* was complemented with either two main (H1.1, H1.2) or all three H1 variants to generate the following lines: *3h1-comp^1,2^*= *h1.1h1.2h1.3;H1.1-RFP;H1.2-GFP* (line #KR276), *3h1-comp^1,2,3^* = *h1.1h1.2h1.3;prom.H1.1::H1.1-RFP;prom.H1.2::H1.2-CFP;*prom.H1.3::H1.3-GFP (lines #KR264 and #KR265). For FRAP experiments the UBQ10::H2B-RFP (Lucas, Kenobi et al. 2013) was crossed with *3h1* and in the subsequent generations by genotyping the *3h1*/UBQ10::H2B-RFP and WT segregants were identified.

Seeds were surface sterilized and rinsed in sterile water before transfer onto germination medium (0.5 x MS medium, 0.8% agar). They were placed on the medium using toothpicks to ensure uniform distribution, stratified 2-4 days at 4°C, and transferred into a plant growth incubator (Percival, Germany) with long-day photoperiod (16 h, 22 °C day/8 h, 18 °C night) and light flux around 120 μM*s^−1^*m^−2^ for routine experiments. Growth of calli and scoring of lateral root production was testing under continuous light (light flux around 100 μM*s^−1^*m^−2^, Aralab FitoClima 1200). When the flowering stage was necessary, the 10 days-old seedlings were transferred into the soil and grown at 19-21°C with a 16h day/8h night photoperiod.

### Chromatin analyses and immunostaining

Nuclei area, heterochromatin (RHF, CCs) and immunostaining analyses were carried out essentially as described (Pavlova, Tessadori et al. 2010) with minor modifications. Nuclei were isolated from rosette leaves of 3-4 weeks old seedlings; per extraction 5 leaves were fixed during 20 min under vacuum in a fresh 4% formaldehyde solution prior to isolation and resuspension of nuclei in a final volume of 1 mL Nuclei Isolation Buffer (NIB). DAPI was added at a concentration of 0.1 mg/ml for flow-sorting according to DNA content. Diploid (2C) nuclei have been flow-sorted using a BD FACSAria IIIu flow cytometer with a 450/50 nm filter (405 nm laser), equipped with a 100-μm nozzle and 25 Psi pressure. Nuclei were collected in 200 μl of NIB before spreading on Superfrost plus slides (1000 nuclei per slide) and stored at 4°C until use. Mutant and wild-type plants were grown and processed for nuclei isolation and immunostaining in parallel.

For heterochromatin analysis, slides were rinsed in SSC2X then PBS before staining with DAPI 1 μg/ml in Vectashield (Vector Laboratory). For immunostaining, the protocol essentially followed previously described steps (Pavlova, Tessadori et al. 2010). As primary antibodies, rabbit anti-Histone H3 (Abcam; ab1791), anti-Histone H1 (Agrisera; as111801), anti-H3K27me3 (Active Motif; 39155), anti-H3K27me1 (Abcam; ab113671), anti-H3K4me3 (Abcam; ab8580), anti-H3K9ac (Abcam; ab10812) and anti-H3K9me1 (Abcam8896) were used at a dilution of 1:200 and incubated at 37°C for 1 h. As secondary antibody, Alexa Fluor 488-conjugated goat anti-rabbit IgG (Molecular Probes; A-11008) was used at a dilution of 1:1000 and incubated for 2 h at 37°C. Nuclei were counterstained for DNA with Propidium Iodide (PI).16-bit images were acquired using a Leica TCS SP5 Confocal Laser Scanning Microscope (CLSM) (Leica microsystems, GmBH, Germany) using a 63× GLY lens (NA 1.4) for heterochromatin and immunostaining analyses. Exposure times, illumination intensities, zoom factor, scanning speed and pinhole were kept identical for the image series in an experiment. For RHF measurements, signal intensities were recorded in manually drawn ROIs capturing chromocenters and normalized over the whole nucleus intensity using Fiji (Schindelin, Arganda-Carreras et al. 2012). For immunostaining, signal intensities for antibodies were normalized against (PI) levels. Graphs were plotted in excel and the data were statistically assessed using a student t-test (unpaired, unequal variance) for comparing wild-type and mutant samples.

### Fluorescent In Situ Hybridization (FISH) and 3D image processing

#### FISH analysis of leaf nuclei

Nuclei were isolated from leaves of 35 days old rosettes grown under a 16h/8h photoperiod. Nuclei extraction and embedding in acrylamide gel pads on slide was done as described (Ashenafi and Baroux 2018). Centromeric and 45S rDNA repeats were detected by FISH using pAL1 and pTA9 to generate DNA probes, respectively (Fransz, De Jong et al. 2002). FISH was done as described (Ashenafi and Baroux 2018) with the following labelling kits and fluorescent immunolabeling reagents: DIG-Nick (Sigma Aldrich, 11745816910), mouse IgG anti-DIG (1:250, Sigma Aldrich, 11333062910), goat IgG anti-mouse IgG~Alexa 488 (1:200, Life Technologies, A-11001); Biotin-Nick translation kit (Sigma Aldrich,11745824910), Biotinylated Anti-Avidin D (1:250, Vector Labs, BA-0300) and Texas Red Avidin D (1:1000, Vector Labs, A-2006). Nuclei were counterstained for DNA with DAPI in Vectashield (Vector Laboratory). FISH signals in 3D nuclei were imaged using STimulated Emission Depletion (STED) microscopy (Leica SP8R WL 3xSTED, Leica microsystems, Germany).

#### FISH analysis of cotyledon nuclei

Nuclei were isolated from dissected cotyledons of 5-day-old seedlings grown under a 16h/8h photoperiod. Nuclei extraction, fixation and hybridization with pAL1-derived and F28D6-derived (180bp-repeats) probes (Fransz, De Jong et al. 2002) was performed as previously described (Bourbousse, Mestiri et al. 2015). Slides were washed and mounted in Vectashield with 2 μg/μL DAPI and image acquisition was performed as in (Bourbousse, Mestiri et al. 2015).

### Nuclei isolation for MNase-seq

Nuclei were isolated from 3 week old seedlings frozen in liquid nitrogen as previously described (Chodavarapu, Feng et al. 2010) with following modifications: after resuspending in HBB, nuclei were applied to layer of HBB with 40% percoll (GE Healthcare), centrifuged at 1000g, 6 min, resuspended in HBB, applied to 40/75% percoll gradient, centrifuged at 400g, 40 min, collected and washed three times with HBC. The integrity of extracted nuclei was monitored using DAPI staining and fluorescence microscopy. The quantity of nuclei was measured by qPCR with primers targeting nuclear DNA.

Digestion was performed by incubating nuclei suspended in DB buffer (16 mM Tris-HCl pH=7.6, 50 mM NaCl, 2.5 mM CaCl_2_, 0.01 mM PMSF, 1× Complete EDTA-free Protease Inhibitors (Roche)) with 1.5 μl (final concentration - 0.3 U/μl) of micrococcal nuclease (Thermo Fisher) and 2 μl (final concentration - 0.2 U/μl) of RNase A (Thermo Fisher) at 8°C for 90 min with gentle mixing. The reaction was stopped by adding equal volume of 2x Lysis buffer with EDTA (100 mM Tris-HCl pH=8, 200 mM NaCl, 50 mM EDTA, 1% SDS). The samples were lysed by incubation in 37°C for 60 min with shaking (1000 rpm). DNA was purified using phenol-chloroform extraction, precipitated with isopropanol and sodium acetate and resuspended in water.

DNA was size selected by electrophoresis on 2% agarose gel with 1xTAE buffer with SYBR Gold (Invitrogen) stain. Mononucleosomal band was excised, frozen and squeezed by three cycles of spinning and rehydration on centrifuge column. DNA was purified and concentrated using Agencourt AMPure XP beads (Beckman Coulter). Barcoded libraries were synthesized from 100 ng of mononucleosomal DNA using Ion Xpress™ Plus gDNA Fragment Library Preparation Kit and Ion Xpress™ Barcode Adapters. DNA was end repaired prior to adapter ligation and size selection and amplification steps were omitted. Resulting libraries were quantified with Ion Library Quantitation Kit, pooled and used to prepare template by clonal PCR with Ion PI™ Template OT2 200 Kit v3 on Ion OneTouch™ 2 System. Sequencing was performed on Ion PI™ chip v2 and Ion Proton™ sequencer using Ion PI™ Sequencing 200 Kit v2 (all Ion Torrent kits and software are trademarks of Thermo Fisher).

### FRAP imaging and data analyses

A *promUBQ10::H2B-RFP* marker (Maizel, von Wangenheim et al. 2011) was introgressed in *3h1* mutant plants by crossing. Both wild-type and triple mutant segregants were analysed. Measurements were done on root tips of two weeks-old seedlings grown as previously described. One sample was prepared at a time: the root was excised and delicately mounted (i.e. without squashing) in 0.5 x MS between slide and coverslip (precleaned with EtOH), sealed with transparent nail polish and let 10 min to equilibrate upside down on the microscope platform before measurements. The imaging chamber was set at a constant temperature of 20°C (higher/fluctuating temperatures induce nuclei juggling). Bleaching and imaging was done using an APO PL 40x oil immersion objective, NA 1.3, over single plane capturing an optical section of ~2μm encompassing a single nucleus (pinhole opening to 5AU) with a 256×256 pixels image format, 3-fold zoom factor. Bleaching was performed in euchromatin within ROI of 1 μm diameter using 5 or more pulses until near total bleach was obtained (Argon laser at 80% power, 100% transmission in 488nm) and post-bleach images were recorded using with 5-7% laser transmission for excitation, in a series of 10 time points, 1 sec interval, followed by 10 time points, 60 sec interval. For analyzing fluorescence recovery, images were first corrected for nuclear drifts occurring during acquisition, using a rigid registration approach in Fiji ((Schindelin, Arganda-Carreras et al. 2012), plugin/registration/stack reg/rigid transformation). When a single image captured several nuclei, single nuclei were cropped for registration and analysis. Fluorescence measurements were done on the bleach ROI, a control ROI near and outside the nucleus, and over the whole nucleus. Calculation of fluorescence recovery was done as described in (Phair, Gorski et al. 2004, Rosa, Ntoukakis et al. 2014) whereby the initial intensity was normalized at 1 for each image before average calculation.

### TEM sample preparation, imaging and image analysis

70 nm tissue sections were prepared from 2 weeks old seedling roots, using a high-pressure freezing/freeze substitution and uranyl acetate staining approach as described in details previously (Fabrice T, Cherkezeyan et al. 2017). Sections from the elongation zone were selected for the analysis (i.e. meristematic zone was avoided) and nuclei pictures were consistently recorded from the epidermal layer at the 24’500 fold magnification yielding a resolution of 1 pixel=2 nm in our setup. For the analysis, square regions of interests (ROI) of similar size (ca 800×800 +/-200 pixels) were captured in euchromatin regions (i.e. excluding strongly staining chromocenters) for the analysis. Spatial autocorrelation analysis delivers a mathematical model of chromatin density distribution for each ROI with respect to the physical length scales within which signal patterns (i.e. local objects of similar intensities) are repeated in a regular pattern (periodicity) (Cherkezyan, Stypula-Cyrus et al. 2014). We used a user-friendly graphical interface developed in Matlab for batch processing of multiple ROIs available at www.github.org\barouxlab\ChromDensityNano and described in details previously (Fabrice T, Cherkezeyan et al. 2017).

### Analysis of developmental transitions

#### Flowering time

Plants for flowering experiments were grown in the greenhouse or growth chamber under the long day light regime. To avoid positional effect, different genotypes were always randomly arranged over growth area. The number of rosette leaves was counted when the inflorescence was about 0.5 cm long.

#### Root length and lateral root scoring

Seedlings were grown vertically on square petri dishes under continuous light regime. The plates were scanned 8 days after germination to score for the number of lateral roots. Root (main and lateral) lengths were scored using manual vector tracing in Fiji, reported at scale (Schindelin, Arganda-Carreras et al. 2012). For microscopic observations of lateral root primordia, five days old seedlings grown under continuous light were fixed in 70% ethanol, rinsed once in sterile water and mounted in water on microscope slides (5 roots aligned/slide covered with 40×22mm coverglass). Primordia were scored according to published developmental scale (Malamy and Benfey 1997). Graphs were plotted in R.

#### Stomata patterning

Fresh epidermal peals of 14 days old cotyledons were mounted in water. Images of the adaxial surface were recorded with DIC microscopy and stomatal clusters were scored following as described (Kutter, Schob et al. 2007).

#### Seed dormancy

The experiment was designed as described previously (Nakabayashi, Bartsch et al. 2012) with minor modifications. Plants were grown in a growth chamber under long day light regime (at least three plants for each genotype) with controlled humidity. Freshly harvested seeds were stored under constant conditions. Around 180 seeds, collected from single plants, were placed on wet filter paper in a Petri dish and incubated in the growth incubator at 22°C under long day light regime. After three days the number of emerging radicles was counted. For the time point “day 1”, seeds one day after harvesting were used. For the time point “3 weeks”, seeds from the same batch were used three weeks after harvesting.

#### Callus induction

Cotyledons from 7 days-old seedlings grown under a 16h/8h photoperiod were excised, transferred onto callus induction medium (CIM, Gamborg B5, 0.05% MES, 2% Glucose, 0.1 mg/L kinetin, 0.5 mg/L 2,4-D) and let to develop for 5 weeks under a 16h/8h photoperiod. Callus size (area) was determined from images using manually drawn contours in Fiji (Schindelin, Arganda-Carreras et al. 2012). Graphs were plotted in R.

### Immunoblot Analyses

Seeds were surface sterilized in 70% ethanol 0.05% SDS for 3 min and rinsed into 90% ethanol before drying and plating on MS medium supplemented with 0.5% sucrose and 0.9% agar. Eight-day-old seedlings were used for chromatin extraction protocol as described previously (Bowler, Benvenuto et al. 2004). Forty micrograms of protein samples, as estimated by the bicinchoninic acid method, were loaded on 14% LiDs Tris-Tricine gels and blotted onto Immobilon-P membranes (Millipore) before immunodetection using Merck Millipore antibodies recognizing either unmodified histone H4 (#05-858), H3K27me3 (#07-360) or custom-made rice histone H2B antibody generated by Prof. David Spiker (Bourbousse, Ahmed et al. 2012).

### RNA-seq

RNA was isolated using modified TRIzol method (Chomczynski and Sacchi 1987). Ribosomal RNA was removed using RiboMinus Plant Kit (Thermo Fisher) and ERCC RNA Spike-In Mix 1 (Thermo Fisher) was added. Libraries were prepaired with Ion Total RNA-Seq Kit v2 and Ion Xpress RNA-Seq Barcode 1-16 Kit according to user guide. Sequencing template was generated with Ion PI™ Template OT2 200 Kit v3 on Ion OneTouch™ 2 System. Sequencing was performed on Ion PI™ chip v2 and Ion Proton™ sequencer using Ion PI™ Sequencing 200 Kit v2 (all Ion Torrent kits and software are trademarks of Thermo Fisher).

Base calling and adapter trimming was performed automatically by Torrent Suite software. Residual rRNA and ERCC reads were identified and filtered out using bbsplit and filterbyname scripts from BBTools suite (Brian Bushnell). Reads were aligned to TAIR10 genome using TMAP 5.0.13. with soft clipping from both ends and returning all the mappings with the best score. Other settings were set according to Torrent Suite defaults. Unaligned reads were aligned with BBMap (Brian Bushnell). Quantitation to ARAPORT11 transcripts and differential expression analysis was performed in Partek Flow (Partek Inc.) using Partek GSA algorithm.

The distribution of genes up-regulated in *3h1* versus all genes in *Arabidopsis thaliana* genome across chromatin states. Genomic coordinates for chromatin states (CS) locations across the genome were downloaded from published data (Sequeira-Mendes, Araguez et al. 2014). The genomic coordinates for genes up-regulated in *3h1* were taken from TAIR 9 to be consistent with CS coordinates. Then, for each gene the percentage of overlapped chromatin states was calculated and for the final graph the summary of all analyzed genes was presented.

### MNase-seq data analysis

Base calling and adapter trimming was performed automatically by Torrent Suite Software. Reads were aligned with TMAP 5.0.13. Soft clipping was turned off, end repair was allowed and all alignments for multi-mapping reads were reported. Other settings were set according to Torrent Suite defaults. Multi mapping read positions were resolved using MMR (Kahles, Behr et al. 2016) with default settings.Peak calling was performed on reads reaching terminal adapter with length range between 147 and 220 nt using set of custom made python scripts. First read centers were piled up and then highest coverage positions were selected using greedy algorithm. Ends of the longest read used to define position were used as peak boundaries. Peak was called only if its boundaries were not overlapping those of neighboring peak. NRLs were defined by calculating peak-to-peak distances from peak calling results. The frequency of distances was calculated as a percentage of all measurements and binned into three groups. Histograms were plotted in Microsoft Excel.

Quantile normalized wiggle occupancy files were generated with DANPOS2 (Chen, Xi et al. 2013) with default settings. To avoid shifting of read positions (automatic procedure for single-end reads), program was fed with “fake 75 nt pared-end” bed files, generated from both ends of alignments of fully-sequenced reads. Wig files were converted to BigWig format using UCSC wigToBigWig (Kent, Zweig et al. 2010) and used in deepTools (Ramirez, Ryan et al. 2016) for plotting.

#### Filtering out Ler residual sequences

Despite series of five backcrosses after introduction of *h1.3* mutant allele from Ler background into our *h1.1h1.2h1.3* (Col-0) line, some residual Ler sequences were still present, mainly neighboring the *H1.3* gene To avoid interference from those sequences in our analyses, we identified their precise genomic coordinates using SNP and coverage analyses by comparing to sequenced genome of parent Ler h1.3 line. We used those coordinates to generate bed files and filter out all reads overlapping residual Ler sequences using bedtools intersect (Quinlan and Hall 2010). Boxplots were obtained with R software.

### MNse-seq and RNA-seq data accession

The data for MNse-seq and RNA-seq discussed in this publication have been deposited in NCBI’s Gene Expression Omnibus (Edgar, Domrachev et al. 2002) and are accessible through GEO Series accession number GSE113558 (https://www.ncbi.nlm.nih.gov/geo/query/acc.cgi?acc=GSE113558). Secure token is qtyrkcsentqrbct.

## Acknowledgements

This work was supported by the Polish National Science Centre (2011/01/N/NZ2/04849) for M.L., the University of Zürich (Forschungskredit) and Plant Fellows Fellowship to K.R., the Swiss National Science Foundation (SNF 310013A/149974 /SystemsX.ch 2014/235 to CB and SNF grants to CR and TNF), Ministry of Sciences and Higher Education grant MNiSW/PO4A/03928 grant for A.J., the Investissements d’Avenir program launched by the French Government and implemented by ANR (ANR-10-LABX-54 MEMOLIFE and ANR-10-IDEX-0001-02 PSL* Research University), and a PhD fellowship from the Université Paris-Sud Doctoral School in Plant Sciences to GT and the COST Action CA1612 INDEPTH. We thank Prof. Ueli Grossniklaus for valuable advice and insightful discussions, for sharing laboratory facilities and support from management personal (V.Gagliardini, C.Eichenberger, A.Bolanos), Daniel Prata for technical assistance in plant growth and selection, Stefan Wyder (URPP, University of Zürich) for advice on statistical tests, D.Bergman (UCLA, USA) for advice on stomata analysis, M. Ingouff (IRD Montpellier, France) for sharing the DynaMET lines, Mariamawit Ashenafi (University of Zurich) for the raw image used for the Figure items 1c-e and technical assistance for immunostaining, Chris Bowler and Vincent Colot (IBENS, Paris) for constant support and helpful discussions to FB, Marek Kalinowski for scripts to analyze chromatin states distribution. We acknowledge the professional service and support of the Cell Sorting and Microscopy Facility of the University of Zürich for nuclei sorting, STED and GSD imaging, TEM preparation, FRAP imaging and advice in data analysis (Urs Ziegler, Jana Dohner, Andres Kaech, Claudia Dumrese and Moritz Kirschman). We thank Marta Koblowska for sharing the facilities of Laboratory of Microarray Analysis for RNA-seq and MNse-seq experiments and Maciej Kotliński for conceptual support in designing and analyzing for RNA-seq and MNse-seq experiments.

## Author contributions

KR, ML, AJ and CB designed the project and the experiments; FB, CR, LC designed and coached specific project parts (cytogenetic and ultrastructural analyses); KR, ML, BM, JS, GT, IM, MK, FT, SF, SG and CB performed experiments/analysed data. KR and CB wrote the manuscript, FB and AJ contributed substantial revisions, all authors read and commented the manuscript.

## Supplemental figure legends

**Figure 1 – figure supplement 1.**
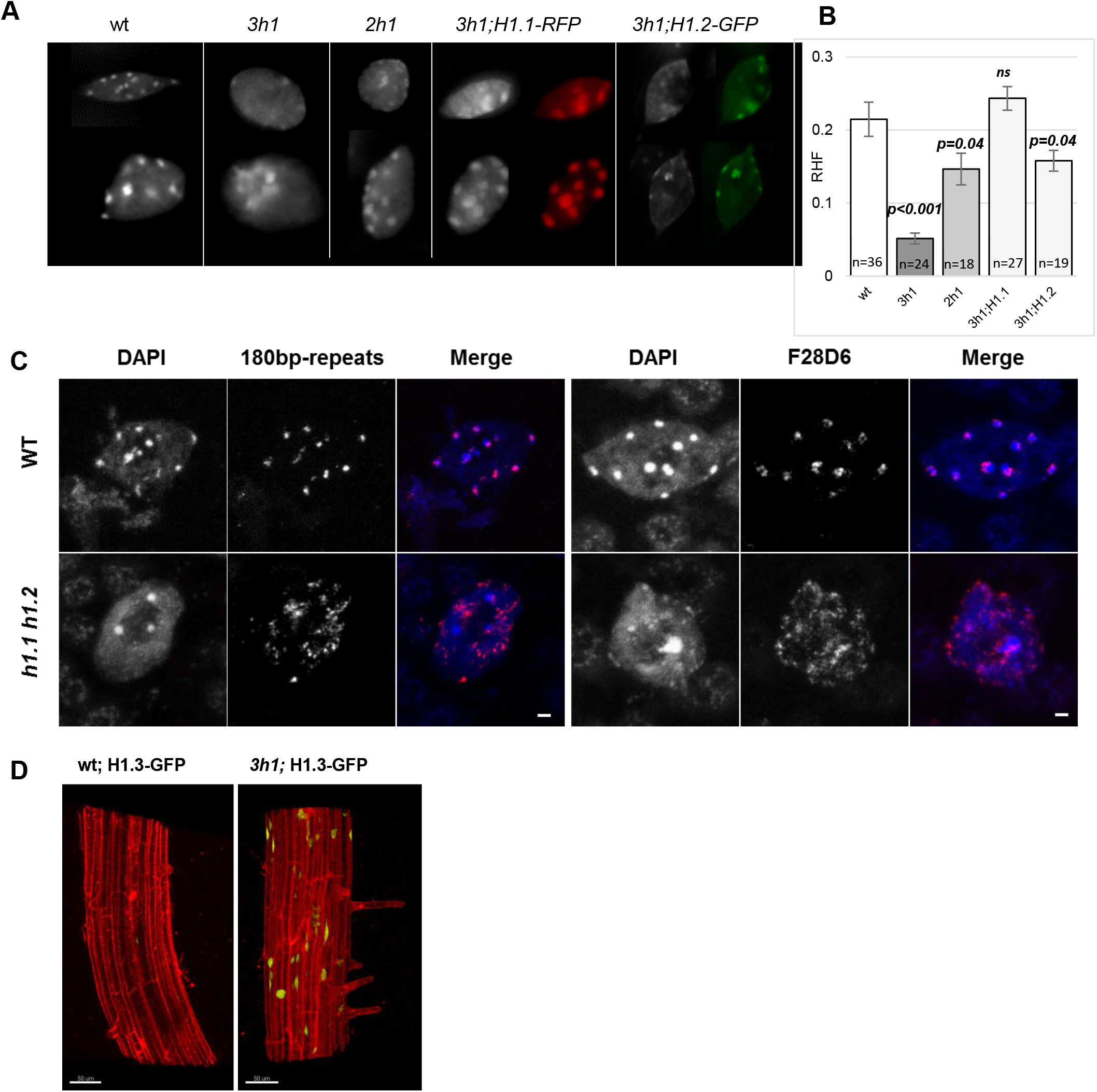
Chromocenter formation relies on H1.1 and H1.2 but not H1.3 in differentiated, adult tissues. (**A**) wt and *h1.1h1.2* nuclei immunostained against centromeric regions with 180bp (centromeric) and F28D6 (pericentromeric) probes and counterstained with DAPI. (**B**) Cytogenetic analyses in root nuclei from triple *h1.1h1.2h1.3* (*3h1*), *double h1.1h1.2* (*2h1*), and triple *3h1* mutants complemented with either H1.1 (*3h1;H1.1-RFP*) or H1.2 (*3h1; H1.2-GFP*) vs wild-type (wt). (**C**) Relative heterochromatin fraction (RHF) in root nuclei is fully or partially restored upon complementation of *3h1* mutant. T-test, error bars, standard error of the mean (s.e.m).

**Figure 1 – figure supplement 2.**
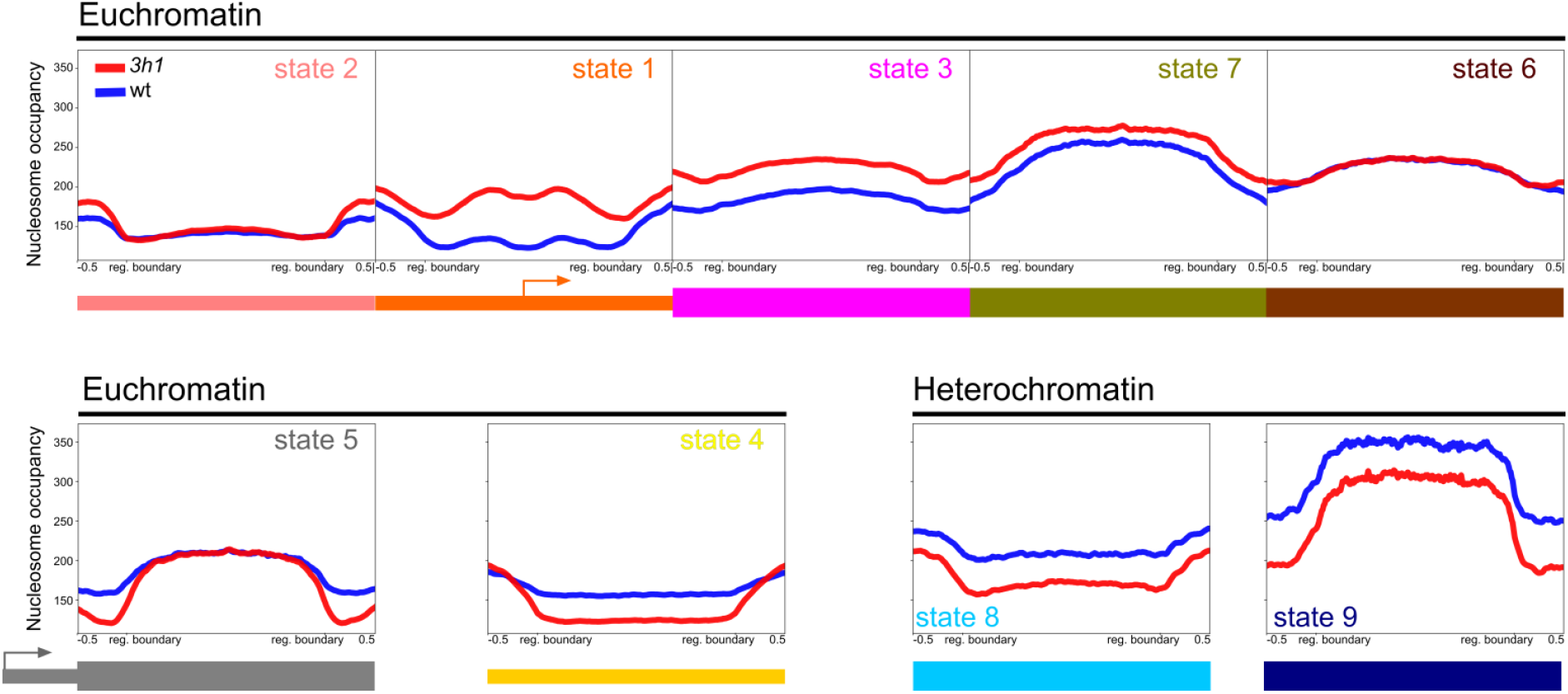
H1 regulates chromatin-state dependent nucleosomal distribution. Nucleosome occupancy per chromatin state (CS) schematically as described by Sequeira-Mendes et al. (Sequeira-Mendes, Araguez et al. 2014) and represented along the most representative genomic feature of each CS as proposed by Vergara and Gutierrez (Vergara and Gutierrez 2017).

**Figure 1 – figure supplement 3.**
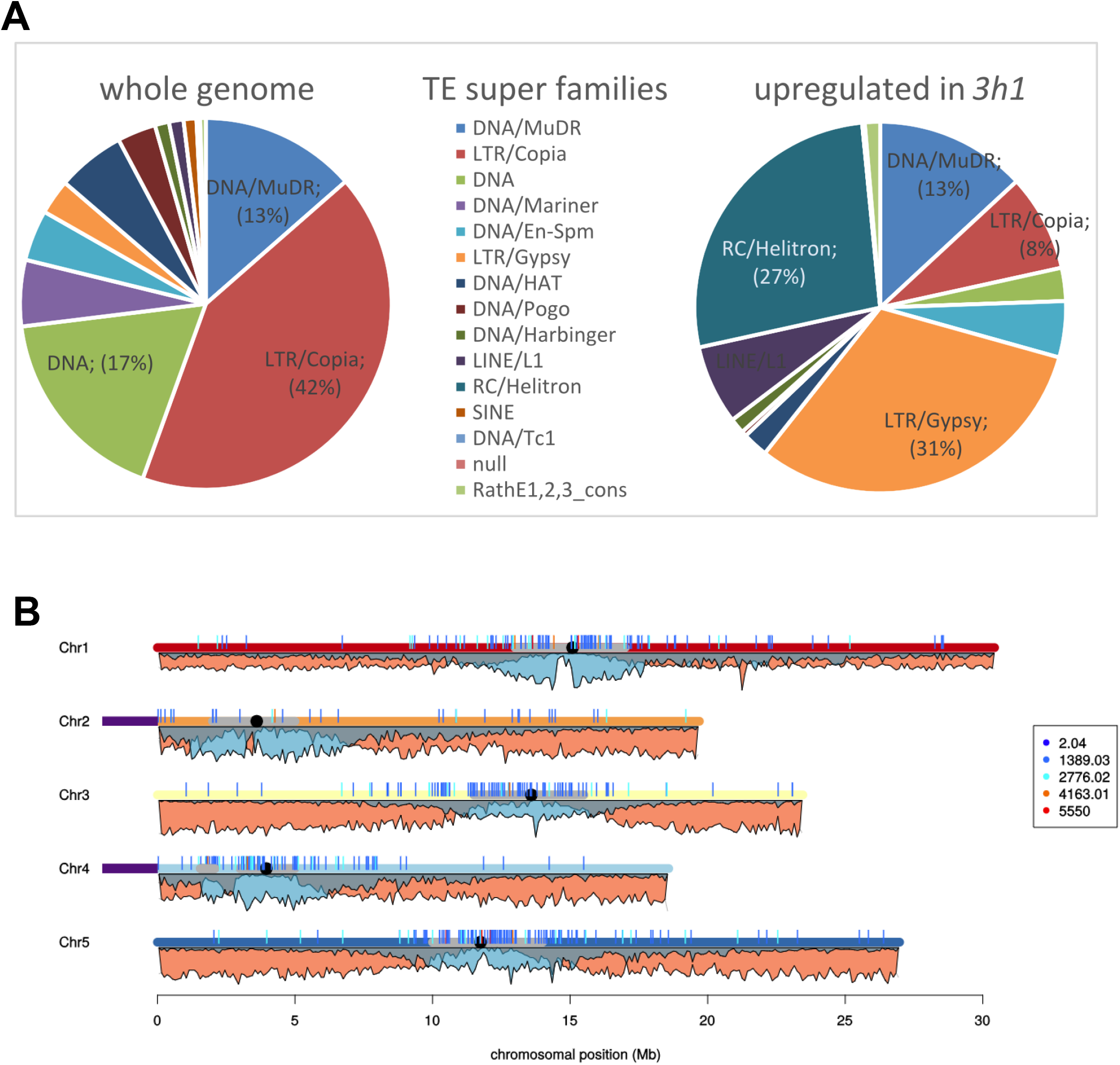
Super families of TEs upregulated in *3h1* mutant. (**A**) The pie charts are based on the data from Table S2. Upregulated elements represent only 1.5% of the TEs, are enriched in Helitron, Copia and Gypsy elements. (**B**) Distribution map of upregulated TEs in *3h1* showing mostly pericentromeric elements. The bars represent single elements, color coded for the fold change expression in *3h1*. The peak-and-valley profiles below each chromosome displays the relative enrichment in genes (orange) and TEs (blue). Graph computed in *R*.

**Figure 2 – figure supplement 1.**
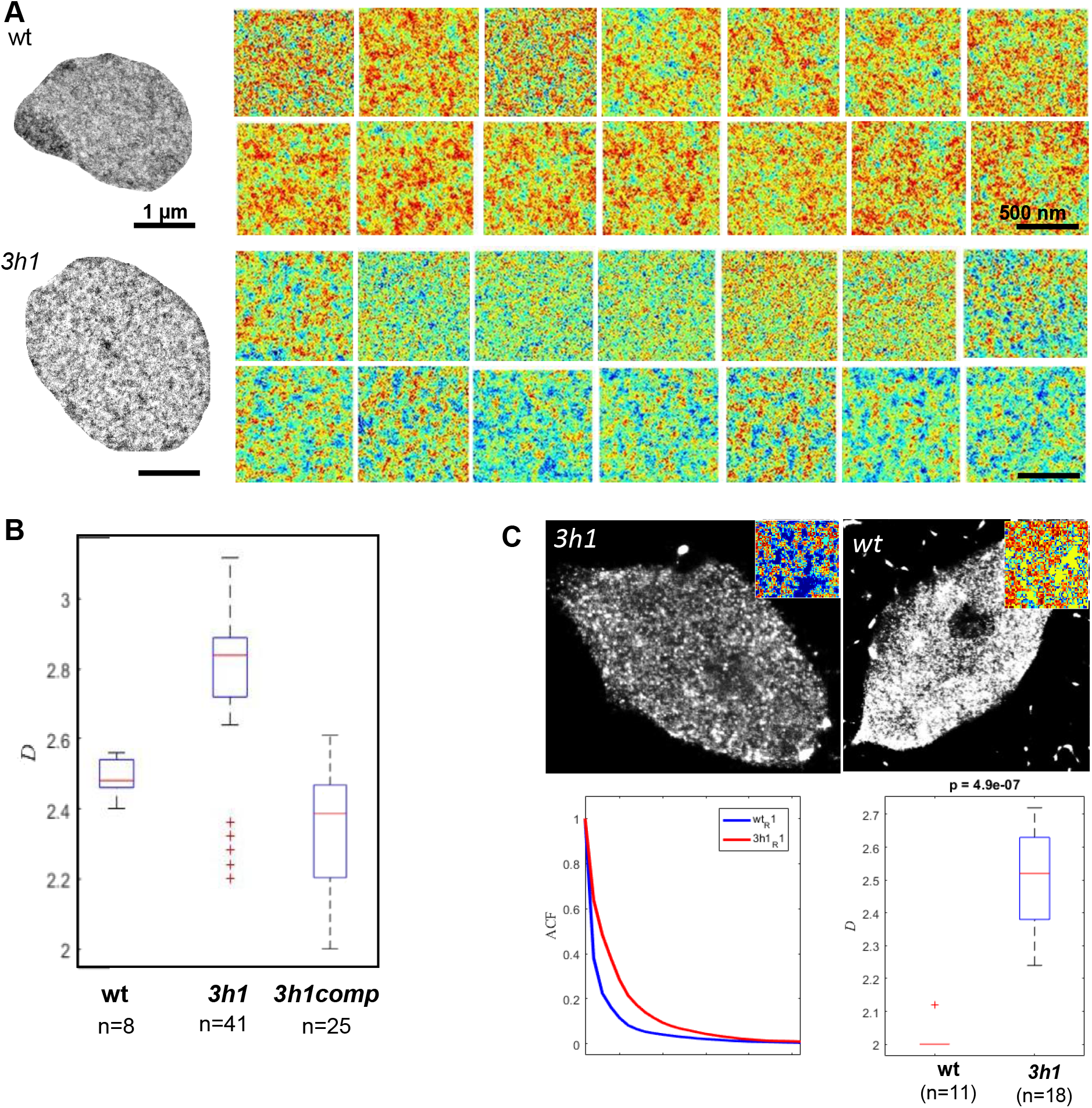
H1-depletion induces spatial dispersion of structural chromatin domains at the nanoscale level. (**A**) Typical TEM image of wild-type (wt) and triple mutant (*3h1*) nuclei (root epidermis) as shown in Figure 2 together with a series of representative regions of interest (ROIs) in euchromatin used for spatial autocorrelation function (ACF) analyses. Scale Bar: 500nm. (**B**) Replicate experiment (TEM sample preparation, imaging and autocorrelation analysis) including a *3h1* mutant line complemented by H1.1-GFP (*3h1comp*) and showing the restoration of a wild-type level of the dispersion (*D*) of length scales in euchromatin. (**C**) The dispersion of nanoscale chromatin domains measured in TEM micrographs is confirmed on super resolution images (GSD imaging) of immunolabelled H3. Analysis as in Figure 2. Inset: ROI as used for ACF analysis.

**Figure 2 – figure supplement 2.**
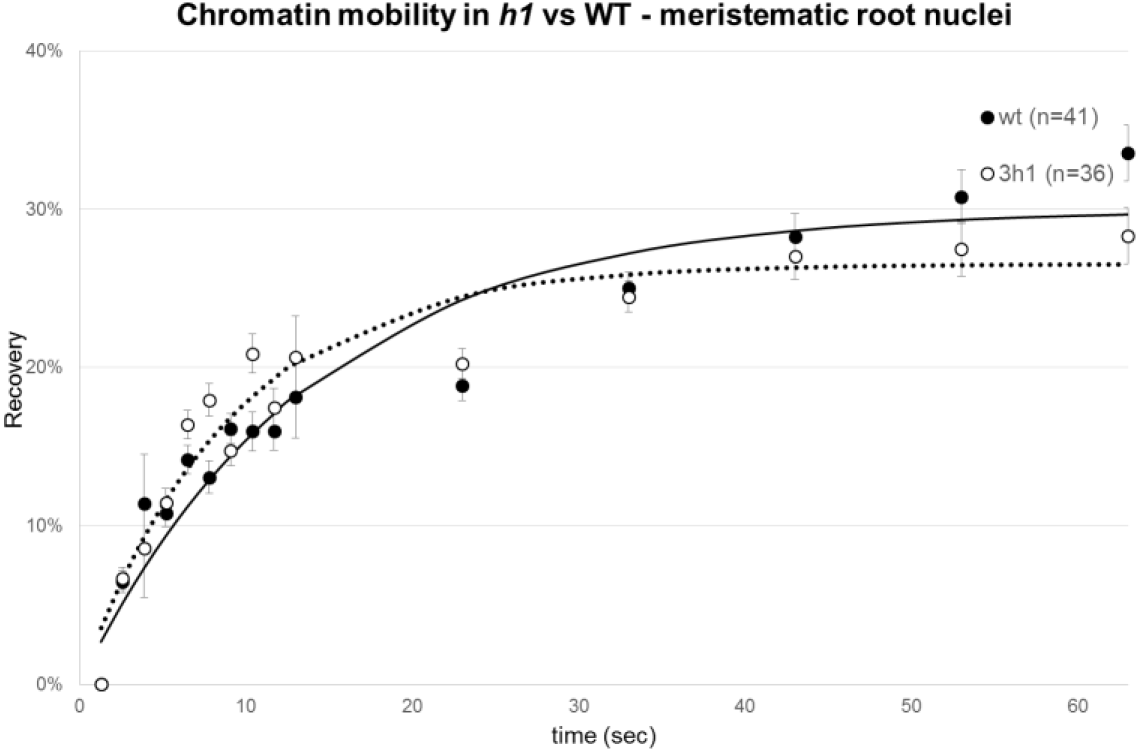
Chromatin mobility in meristematic nuclei is not affected by H1 depletion. Fluorescence recovery after photobleaching (FRAP) in meristematic root nuclei in wt and *3h1* mutant (graph, double normalisation, as done for Figure 2. see Methods). H1 depletion does not alter chromatin mobility in meristematic nuclei. Note that the recovery rate of H1-depleted differentiated nuclei (Figure 2) is similar to that of wt meristematic nuclei.

**Figure 3 – figure supplement 1.**
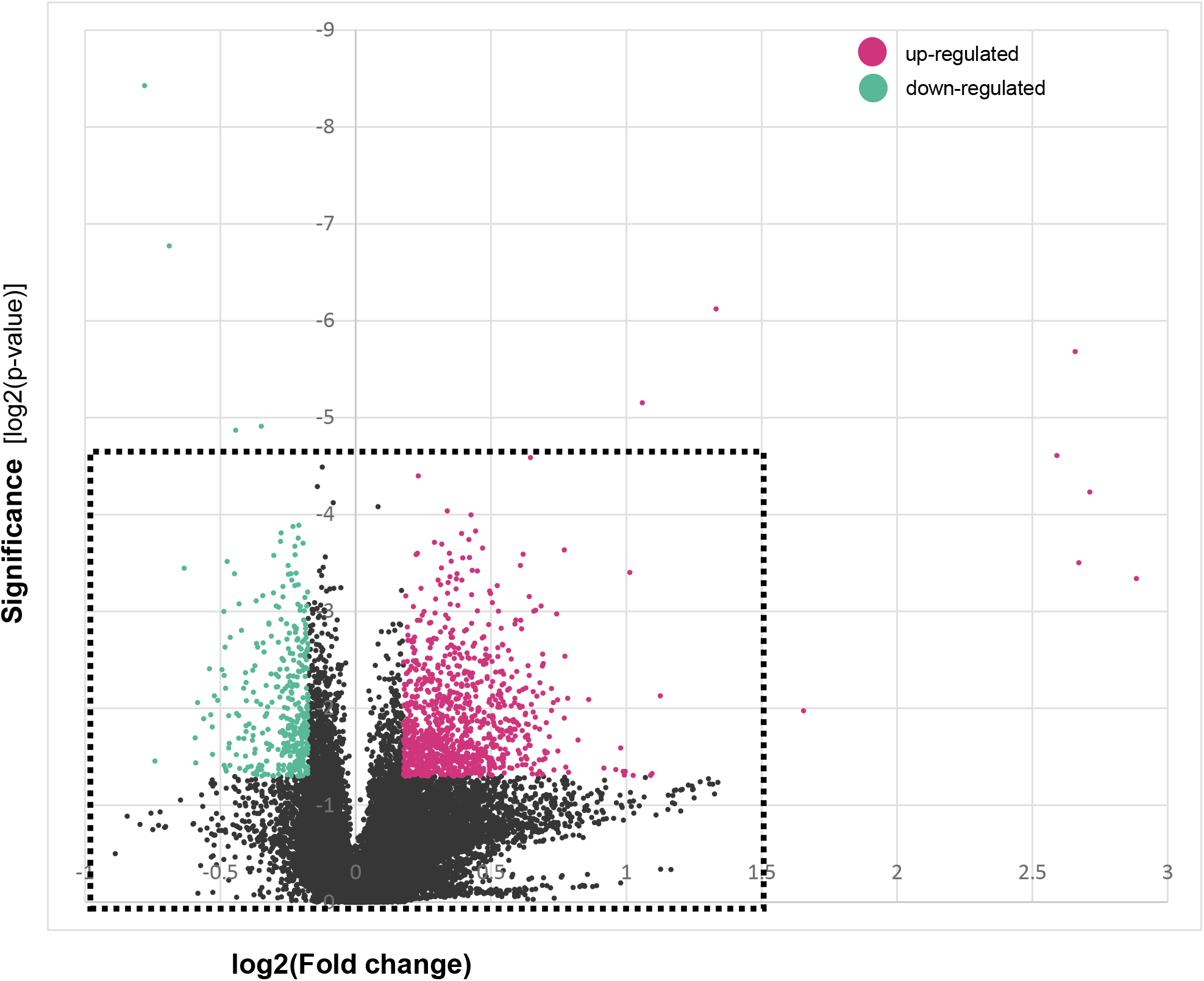
Gene expression in 3h1 mutant. Volcano plot showing significance of gene expression changes in *3h1* vs wt and preferences to up-regulation in *3h1*. This plot is the full version as the one presented Figure 3 (cropped around the dashed box for display purposes).

**Figure 3 – figure supplement 2.**
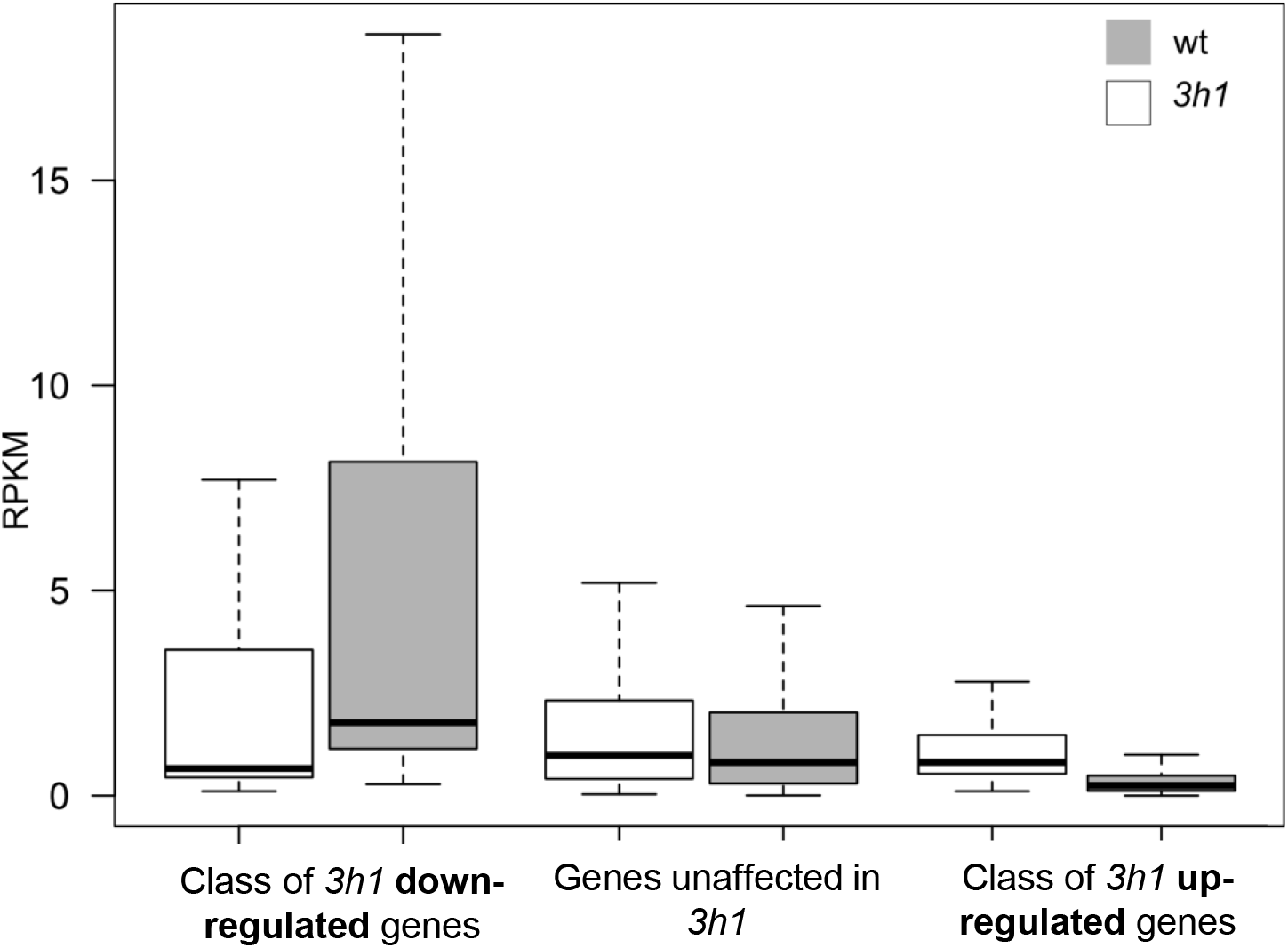
Up‐ and down-regulated genes in *3h1* correspond to gene categories with distinct expression strength in wild-type. The graphs shows the mean expression level in RNAseq profiles for the classes of genes up‐ or down-regulated in *3h1*, or unaffected. The graph shows a clear cut trend in gene classes with respect to their original expression strength in wild-type: *3h1* down-regulated genes represent a class of normally highly expressed genes in wild-type compared to the class of *3h1* up-regulated genes that represent a class of genes with low expression levels in wild-type. Average gene expression (RPKM) from 3 biological replicates for each group of genes: down-regulated in *3h1* (N=43), not-regulated in *3h1* (N=22557) and up-regulated in *3h1* (N=231); p-value ≤ 0.01 and fold change ≥ 1.5.

**Figure 3 – figure supplement 3.**
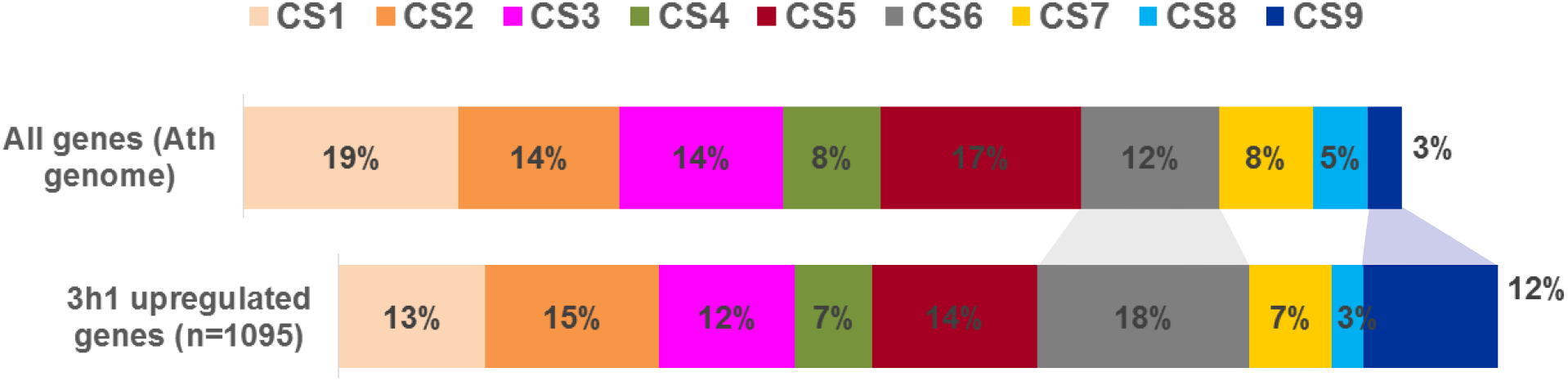
Distribution of chromatin states (CS) of all genes in the *Arabidopsis thaliana* genome versus genes upregulated in *3h1* plants. Among 1095 genes which were up-regulated in *3h1* plants (p-value ≤ 0.05 and fold change ≥ 1.5), 142 are TE genes and 12 are pseudogenes what might explain the slight overrepresentation of CS9. CS6 is also slightly more represented among upregulated genes but otherwise there is no specific category.

**Figure 3 – figure supplement 4.**
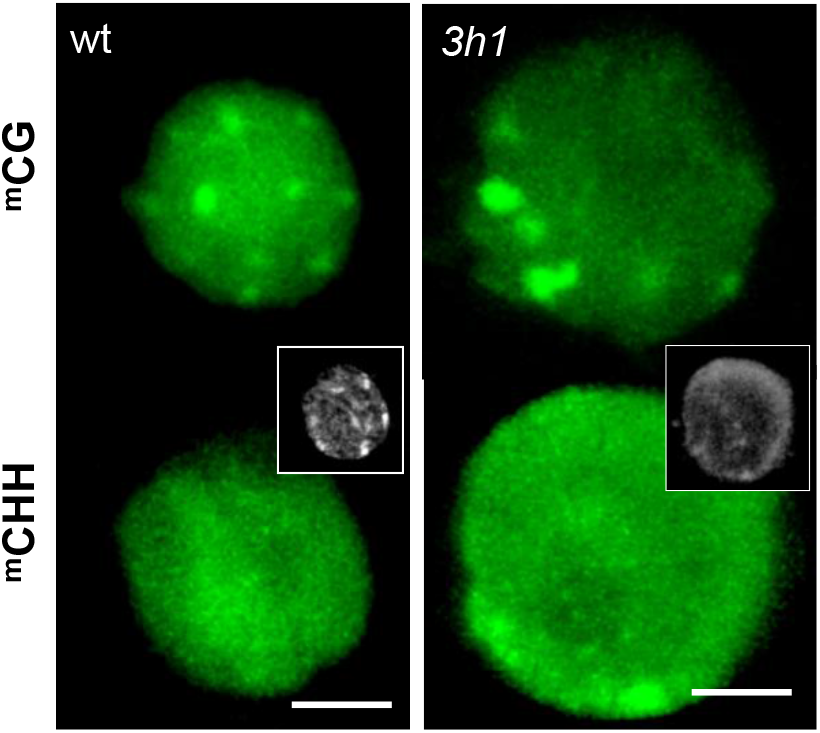
The overall distribution of CHH and CG is not affected in *3h1* mutant nuclei. Confocal imaging of root nuclei in 8 days old seedlings expressing the DynaMET reporters marking methylated DNA in the CG or CHH context as indicated (Ingouff et al, 2017).

**Figure 3 – figure supplement 5.**
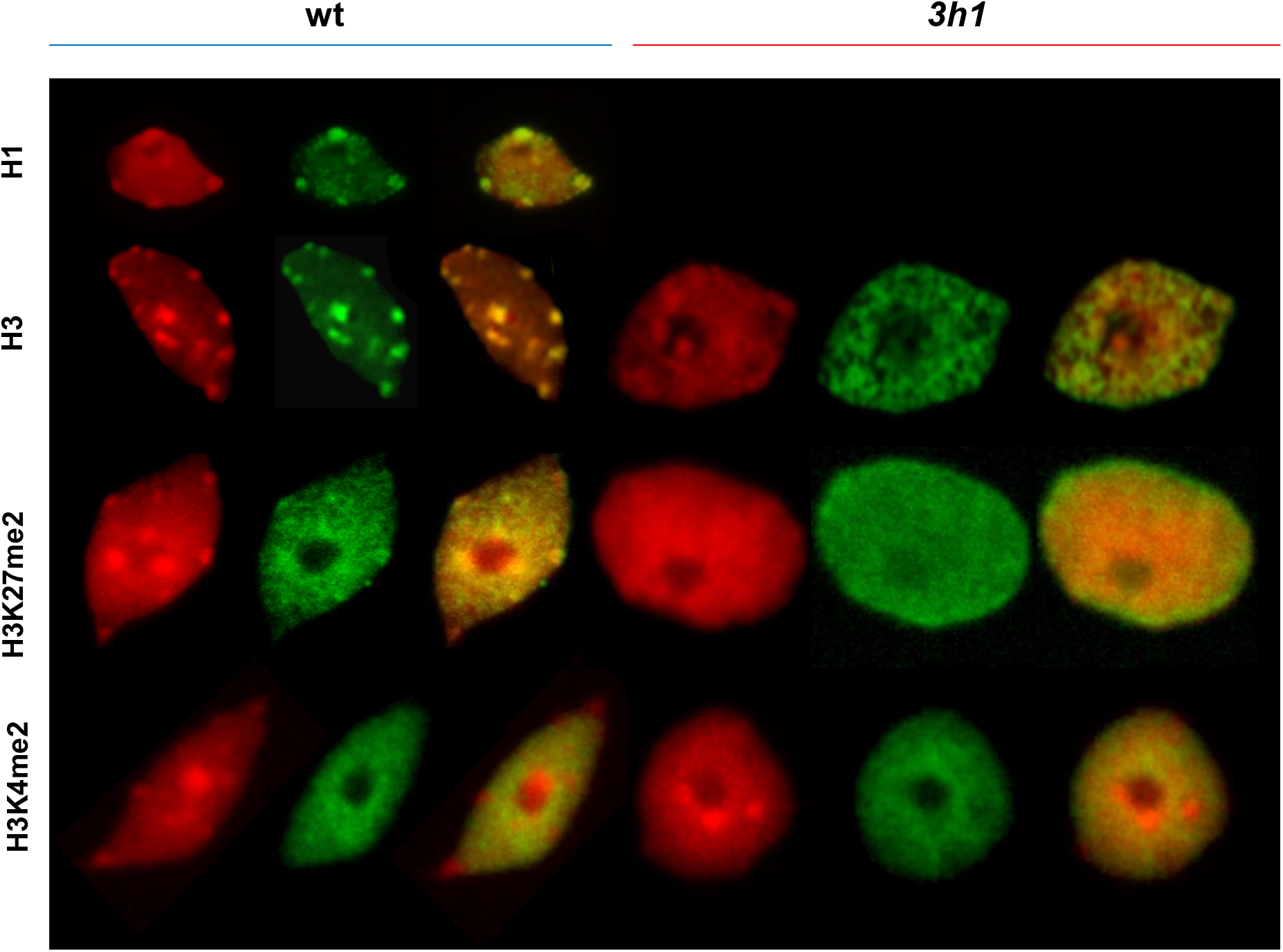
H3, H3K4me2 and H3K27me2 are not affected by H1 depletion. Leaf nuclei isolated, flow-sorted according to their 2C DNA content, spread and immunostained as described in the methods. H1 immunostaining was used as control.

**Figure 3 – figure supplement 6.**
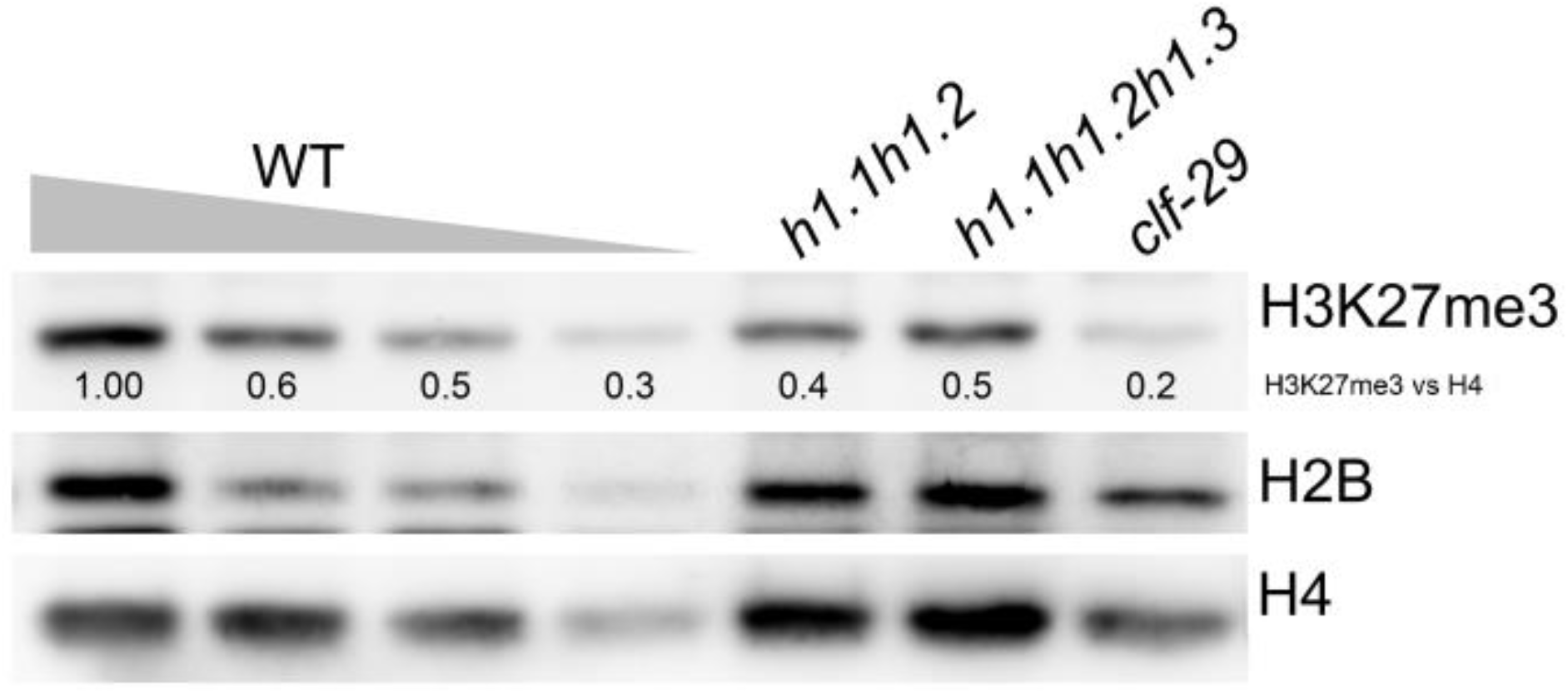
Global levels of H3K27me3 are reduced in *3h1* seedlings. Forty micrograms of chromatin extracts from 8-day-old plants of the different genotypes were analyzed by immunoblot using the indicated antibodies. A dilution series of wild-type plant extracts serves as a quantitative estimation.

**Figure 4 – figure supplement 1.**
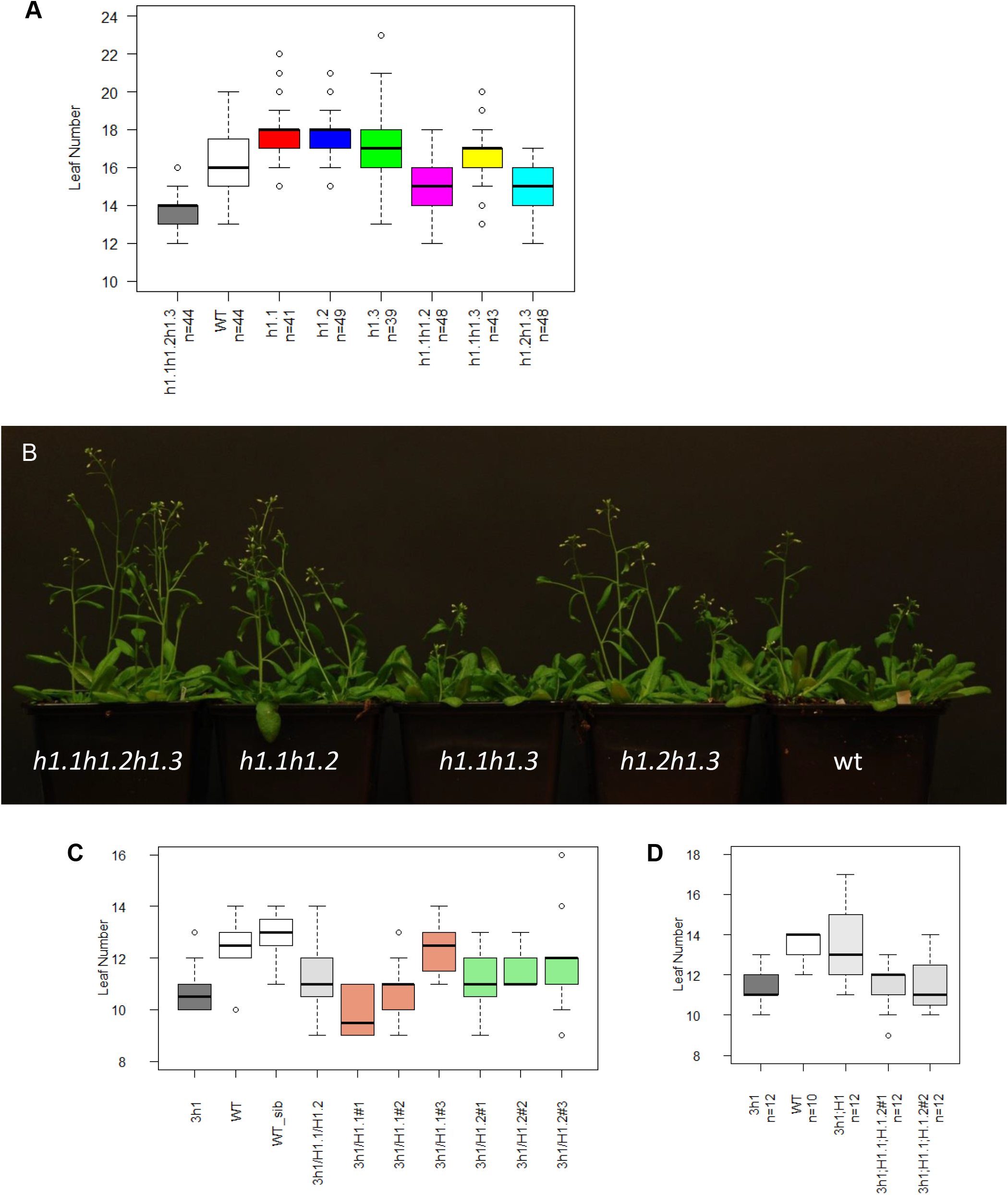
Flowering time for *h1* mutants and complemented lines. (**A**) Leaf number at bolting (~0.5 cm stem) was monitored for different variant combinations of H1 mutants under long day conditions (16h day/8 h night). (**B**) Comparison between wt, triple *h1.1h1.2h1.3* (*3h1*) and three different double *h1.1h1.2, h1.1h1.3, h1.2h1.3* mutants. (**C**) Introducing one or two main H1 variants into *3h1* background does not complement early flowering phenotype. (**D**) Early flowering phenotype in *3h1* was complemented by introducing all three H1 variants (*3h1;H1*). *Legend: 3h1*/H1.1 - *3h1*;prom.H1.1:H1.1-RFP; *3h1*/H1.2 - *3h1*;prom.H1.2::H1.2-GFP; *3h1*/H1.1/H1.2 - *3h1*;prom.H1.1:H1.1-RFP /prom.H1.2::H1.2-GFP; *3h1*;H1 - *3h1*;prom.H1.1:H1.1-RFP /prom.H1.2::H1.2-CFP/prom.H1.3::H1.3-GFP; #1, #2, #3 mean different, independent Arabidopsis lines.

**Figure 4 – figure supplement 2.**
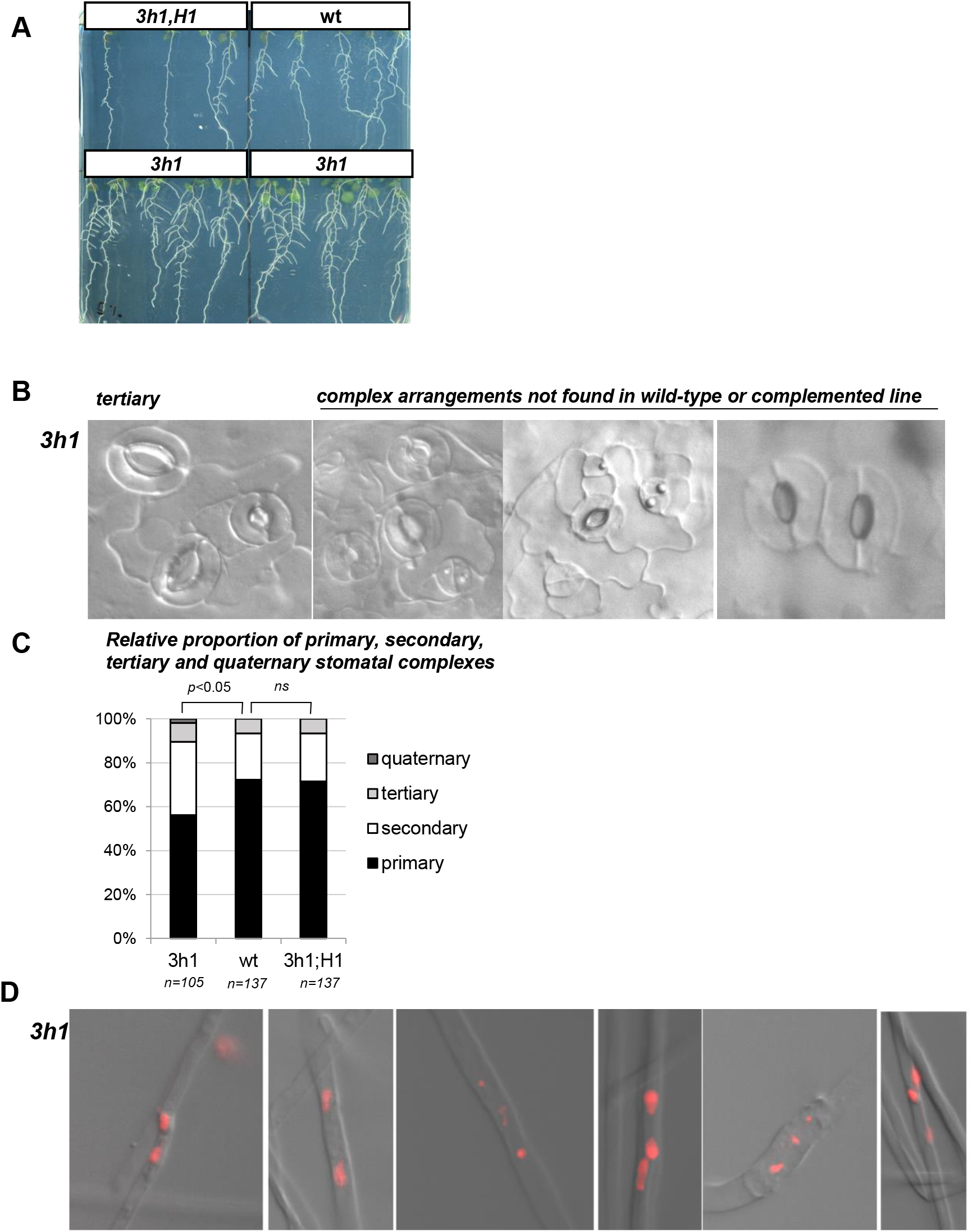
H1 is required for correct developmental transitions. This figure shows additional image and quantification material supporting Figure 4 (**A**) Typical seedling phenotypes in *3h1, 3h1* complemented lines and wt showing differences in lateral root number (**B**) Altered stomata spacing in *3h1* cotyledons (15 days after germination, adaxial side). (**C**) Relative proportion of primary, secondary, tertiary and quaternary stomatal complexes presented in panel B. (**D**) Multicellular root hairs in *3h1* mutant showing ectopic nuclei (DAPI staining, red). Multicellular root hairs were rare in *3h1* (<1% root hairs observed among 3x 24 seedlings) but never observed in wild-type among 3 independent experiments (24 three weeks old seedlings/experiment grown on MS complemented with 1% sugar) *Legend: 3h1*, triple mutant *h1.1;h1.2;h1.3. 3h1;H1*, triple mutant complemented with the three H1 variants tagged with FPs: *3h1;prom.H1.1:H1.1-RFP; prom.H1.2::H1.2-CFP; prom.H1.3::H1.3-GFP.*

**Figure 4 – figure supplement 3.**
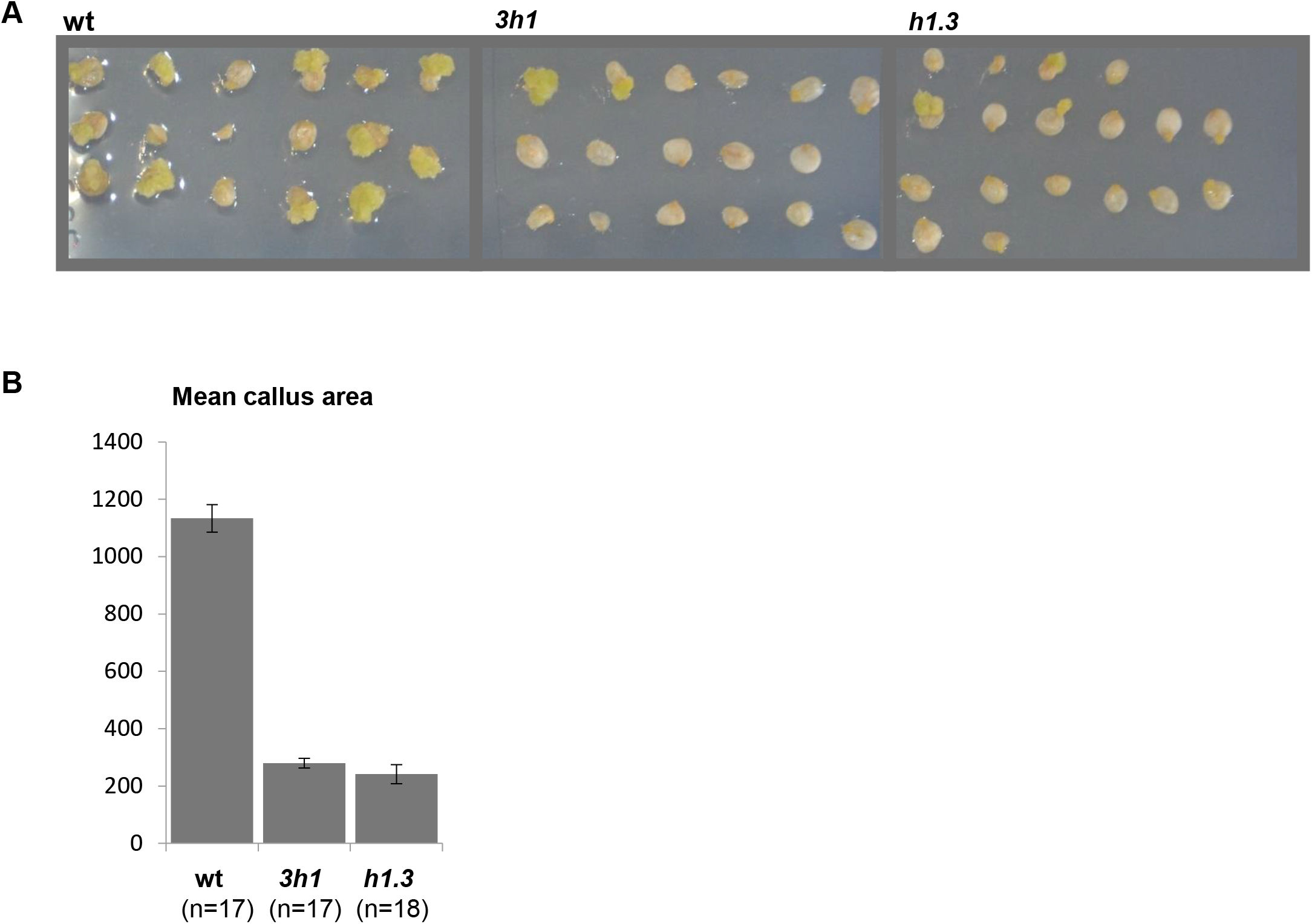
Callus formation efficiency in H1 deficient mutants is reduced. (**A**) Comparison between callus formation in wt, *3h1* and *h1.3.* (**B**) Callus area was measured for wt, *3h1* and *h1.3* with ImageJ.

**Figure 4 – Figure supplement 4.**
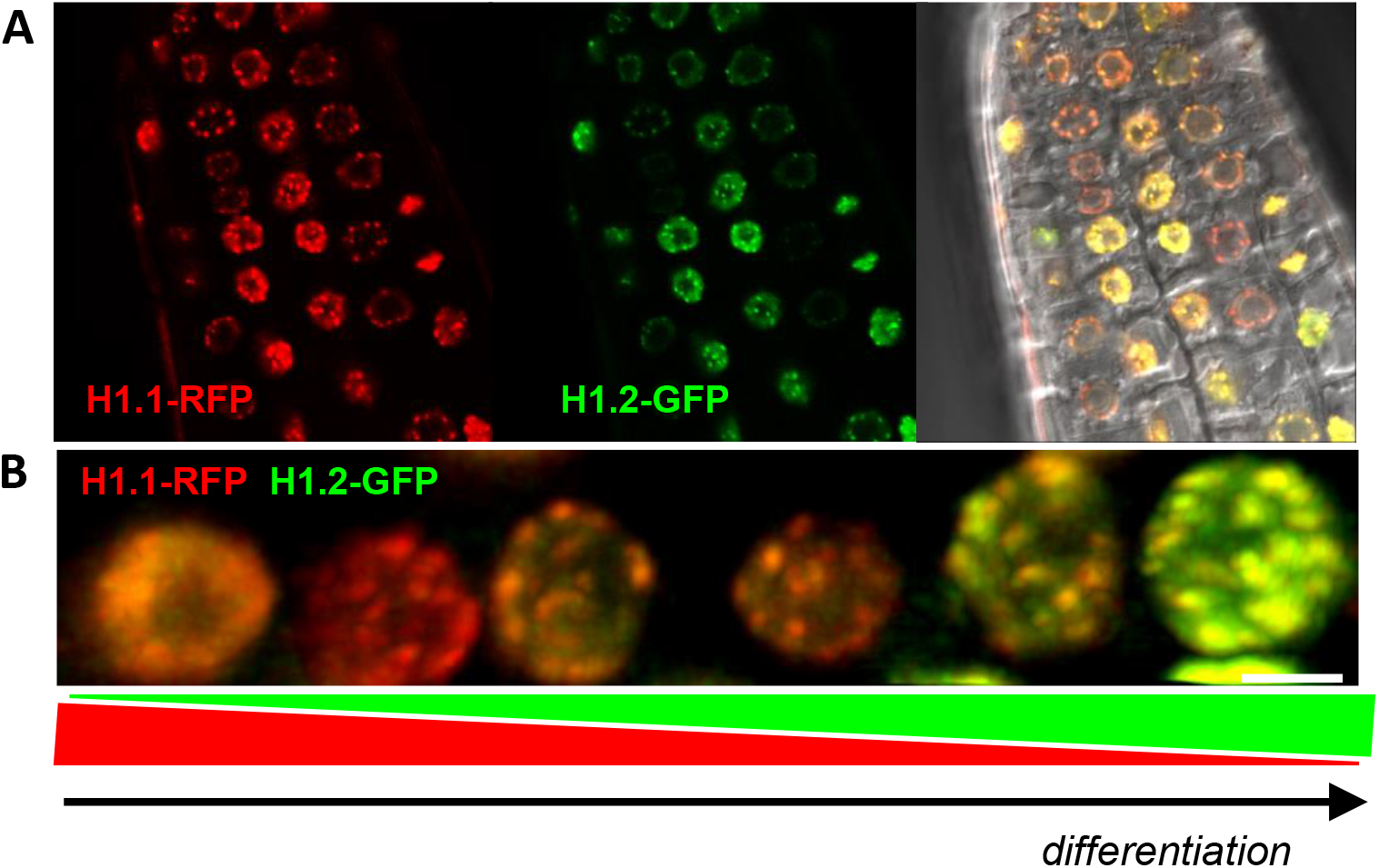
H1.2 levels decrease relative to H1.1 during cellular differentiation in root. (**A**) Snapshot (single plane, CSLM imaging) of a root tip co-expressing H1.1-RFP and H1.2 GFP (as indicated) in the *3h1* background overlaid with the DIC channel (right). (**B**). Representative nuclei series from a single root tip from the meristematic zone to the differentiation zone (3D projection of CSLM series).

## Supplemental tables

**Supplemental Table 1.** Transposable element (TE) expression in *3h1. Available as an Excel table.*

**Supplemental Table 2.** Classes of TEs up-regulated in *3h1*.

| **Super Family** | **Whole genome**<br>**number of TE (%)** |  | **Upregulated in *3h1* (Fch≥2, *P*≤0.05)**<br>**number of TE (%)** |  |
| --- | --- | --- | --- | --- |
| DNA/MuDR | 5410 | (13%) | 59 | (13%) |
| LTR/Copia | 1781 | (42%) | 38 | (8%) |
| DNA | 1829 | (17%) | 13 | (3%) |
| DNA/Mariner | 151 | (6%) | 0 | (0%) |
| DNA/En-Spm | 941 | (4%) | 22 | (5%) |
| LTR/Gypsy | 4181 | (3%) | 141 | (31%) |
| DNA/HAT | 1035 | (6%) | 10 | (2%) |
| DNA/Pogo | 344 | (3%) | 2 | (0%) |
| DNA/Harbinger | 379 | (1%) | 6 | (1%) |
| LINE/L1 | 1447 | (1%) | 31 | (7%) |
| RC/Helitron | 12945 | (0%) | 121 | (27%) |
| SINE | 131 | (1%) | 0 | (0%) |
| DNA/Tc1 | 95 | (0%) | 0 | (0%) |
| null | 16 | (0%) | 1 | (0%) |
| RathE1,2,3_cons | 391 | (0%) | 6 | (1%) |
| **All** | **31076** | **(100%)** | **450** | **(100.0%)** |

**Supplemental Table 3.** Gene expression in *3h1. Available as an Excel table.*

**Supplemental Table 4.** Gene Ontology (GO) analysis of genes which are misregulated in *3h1* mutant.

| **<u>Biological process</u>** |  |  |
| --- | --- | --- |
| **GO Term** | **GO Annotation** | **p-value** |
| photosynthesis, light harvesting in photosystem I | GO:0009768 | 5.59E-20 |
| photosynthesis | GO:0015979 | 2.42E-17 |
| protein-chromophore linkage | GO:0018298 | 1.50E-16 |
| photosynthesis, light harvesting | GO:0009765 | 1.73E-16 |
| photosynthesis, light reaction | GO:0019684 | 7.89E-13 |
| generation of precursor metabolites and energy | GO:0006091 | 5.51E-08 |
| response to light stimulus | GO:0009416 | 6.51E-08 |
| response to radiation | GO:0009314 | 1.62E-07 |
| response to abiotic stimulus | GO:0009628 | 5.20E-07 |
| chlorophyll biosynthetic process | GO:0015995 | 2.14222E-06 |
| porphyrin-containing compound biosynthetic process | GO:0006779 | 5.9281E-06 |
| tetrapyrrole biosynthetic process | GO:0033014 | 8.40871E-06 |
| response to far red light | GO:0010218 | 1.65334E-05 |
| chlorophyll metabolic process | GO:0015994 | 3.82708E-05 |
| response to red light | GO:0010114 | 4.06136E-05 |
| response to high light intensity | GO:0009644 | 0.000107761 |
| porphyrin-containing compound metabolic process | GO:0006778 | 0.000114665 |
| tetrapyrrole metabolic process | GO:0033013 | 0.000119558 |
| response to blue light | GO:0009637 | 0.00021512 |
| response to light intensity | GO:0009642 | 0.000226407 |
| response to stimulus | GO:0050896 | 0.000376309 |
| response to red or far red light | GO:0009639 | 0.001070904 |
| response to low light intensity stimulus | GO:0009645 | 0.005522224 |
| pigment biosynthetic process | GO:0046148 | 0.00574522 |
| protoporphyrinogen IX biosynthetic process | GO:0006782 | 0.006858986 |
| protoporphyrinogen IX metabolic process | GO:0046501 | 0.006858986 |
| response to chemical | GO:0042221 | 0.025952734 |
| pigment metabolic process | GO:0042440 | 0.026767756 |
| heme biosynthetic process | GO:0006783 | 0.04207751 |
| **<u>Molecular function</u>** |  |  |
| **GO Term** | **GO Annotation** | **p-value** |
| pigment binding | GO:0031409 | 6.23E-19 |
| chlorophyll binding | GO:0016168 | 1.43E-16 |
| tetrapyrrole binding | GO:0046906 | 1.4425E-05 |
| oxidoreductase activity | GO:0016628 | 0.025587703 |
| **<u>Cell compartment</u>** |  |  |
| **GO Term** | **GO Annotation** | **p-value** |
| photosystem I | GO:0009522 | 5.61E-27 |
| plastid thylakoid | GO:0031976 | 5.67E-27 |
| chloroplast thylakoid | GO:0009534 | 7.79E-27 |
| thylakoid | GO:0009579 | 2.31E-26 |
| photosystem | GO:0009521 | 1.55E-25 |
| plastid thylakoid membrane | GO:0055035 | 3.20E-23 |
| chloroplast thylakoid membrane | GO:0009535 | 3.41E-23 |
| thylakoid membrane | GO:0042651 | 1.86E-22 |
| photosynthetic membrane | GO:0034357 | 1.97E-22 |
| chloroplast part | GO:0044434 | 3.96E-22 |
| thylakoid part | GO:0044436 | 4.50E-22 |
| organelle subcompartment | GO:0031984 | 4.79E-22 |
| plastid part | GO:0044435 | 1.32E-21 |
| plastoglobule | GO:0010287 | 4.78E-20 |
| light-harvesting complex | GO:0030076 | 7.96E-18 |
| chloroplast envelope | GO:0009941 | 8.15E-18 |
| plastid envelope | GO:0009526 | 2.10E-17 |
| chloroplast stroma | GO:0009570 | 2.24E-14 |
| intracellular organelle part | GO:0044446 | 2.46E-14 |
| organelle part | GO:0044422 | 2.81E-14 |
| photosystem II | GO:0009523 | 4.52E-14 |
| plastid stroma | GO:0009532 | 8.85E-14 |
| plastid | GO:0009536 | 1.60E-13 |
| chloroplast | GO:0009507 | 1.24E-12 |
| envelope | GO:0031975 | 6.97E-12 |
| organelle envelope | GO:0031967 | 7.04E-12 |
| membrane protein complex | GO:0098796 | 1.38E-10 |
| membrane | GO:0016020 | 6.96E-09 |
| cytoplasmic part | GO:0044444 | 1.12E-08 |
| cytoplasm | GO:0005737 | 5.33E-07 |
| macromolecular complex | GO:0032991 | 8.12E-07 |
| photosystem I reaction center | GO:0009538 | 2.05082E-06 |
| protein complex | GO:0043234 | 0.000623038 |
| photosystem II antenna complex | GO:0009783 | 0.00338651 |
| membrane part | GO:0044425 | 0.006706641 |
| integral component of membrane | GO:0016021 | 0.007946083 |
| chloroplast thylakoid membrane protein complex | GO:0098807 | 0.008141248 |
| intracellular ribonucleoprotein complex | GO:0030529 | 0.013895558 |
| ribonucleoprotein complex | GO:1990904 | 0.013895558 |
| intrinsic component of membrane | GO:0031224 | 0.016362585 |
| cell periphery | GO:0071944 | 0.017709796 |
| cell-cell junction | GO:0005911 | 0.019623792 |
| cell junction | GO:0030054 | 0.019623792 |
| cell part | GO:0044464 | 0.019966572 |
| plasmodesma | GO:0009506 | 0.020005906 |
| symplast | GO:0055044 | 0.020005906 |
| cell | GO:0005623 | 0.022080537 |
| chloroplast photosystem II | GO:0030095 | 0.022823861 |
| non-membrane-bounded organelle | GO:0043228 | 0.022958631 |
| intracellular non-membrane-bounded organelle | GO:0043232 | 0.022958631 |
| cytosol | GO:0005829 | 0.023594558 |
| organelle | GO:0043226 | 0.025609296 |
| intracellular organelle | GO:0043229 | 0.025722053 |
| nucleolus | GO:0005730 | 0.026025793 |
| ribosome | GO:0005840 | 0.026106284 |
| cell wall | GO:0005618 | 0.026886069 |
| external encapsulating structure | GO:0030312 | 0.026886069 |
| chloroplast membrane | GO:0031969 | 0.031620021 |
| membrane-bounded organelle | GO:0043227 | 0.036306654 |
| intracellular membrane-bounded organelle | GO:0043231 | 0.03641433 |
| plastid membrane | GO:0042170 | 0.036481998 |
| apoplast | GO:0048046 | 0.041207145 |
| plasma membrane | GO:0005886 | 0.0462473 |
| cytosolic ribosome | GO:0022626 | 0.047006336 |

**Supplemental Table 5.**
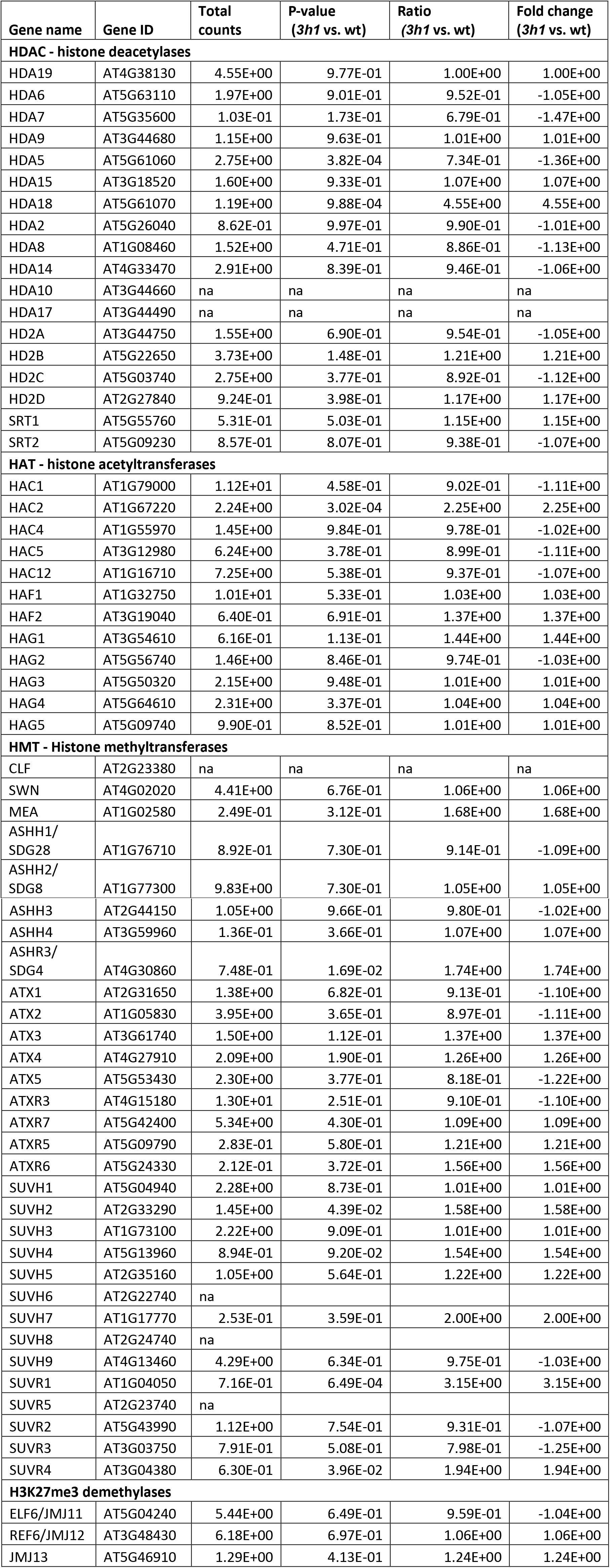
Expression of histone modifying enzymes in *3h1*.

| Gene name | Gene ID | Total counts | P-value (*3h1* vs. wt) | Ratio (*3h1* vs. wt) | Fold change (*3h1* vs. wt) |
| --- | --- | --- | --- | --- | --- |
| **HDAC - histone deacetylases** |  |  |  |  |  |
| HDA19 | AT4G38130 | 4.55E+00 | 9.77E-01 | 1.00E+00 | 1.00E+00 |
| HDA6 | AT5G63110 | 1.97E+00 | 9.01E-01 | 9.52E-01 | -1.05E+00 |
| HDA7 | AT5G35600 | 1.03E-01 | 1.73E-01 | 6.79E-01 | -1.47E+00 |
| HDA9 | AT3G44680 | 1.15E+00 | 9.63E-01 | 1.01E+00 | 1.01E+00 |
| HDA5 | AT5G61060 | 2.75E+00 | 3.82E-04 | 7.34E-01 | -1.36E+00 |
| HDA15 | AT3G18520 | 1.60E+00 | 9.33E-01 | 1.07E+00 | 1.07E+00 |
| HDA18 | AT5G61070 | 1.19E+00 | 9.88E-04 | 4.55E+00 | 4.55E+00 |
| HDA2 | AT5G26040 | 8.62E-01 | 9.97E-01 | 9.90E-01 | -1.01E+00 |
| HDA8 | AT1G08460 | 1.52E+00 | 4.71E-01 | 8.86E-01 | -1.13E+00 |
| HDA14 | AT4G33470 | 2.91E+00 | 8.39E-01 | 9.46E-01 | -1.06E+00 |
| HDA10 | AT3G44660 | na | na | na | na |
| HDA17 | AT3G44490 | na | na | na | na |
| HD2A | AT3G44750 | 1.55E+00 | 6.90E-01 | 9.54E-01 | -1.05E+00 |
| HD2B | AT5G22650 | 3.73E+00 | 1.48E-01 | 1.21E+00 | 1.21E+00 |
| HD2C | AT5G03740 | 2.75E+00 | 3.77E-01 | 8.92E-01 | -1.12E+00 |
| HD2D | AT2G27840 | 9.24E-01 | 3.98E-01 | 1.17E+00 | 1.17E+00 |
| SRT1 | AT5G55760 | 5.31E-01 | 5.03E-01 | 1.15E+00 | 1.15E+00 |
| SRT2 | AT5G09230 | 8.57E-01 | 8.07E-01 | 9.38E-01 | -1.07E+00 |
| **HAT - histone acetyltransferases** |  |  |  |  |  |
| HAC1 | AT1G79000 | 1.12E+01 | 4.58E-01 | 9.02E-01 | -1.11E+00 |
| HAC2 | AT1G67220 | 2.24E+00 | 3.02E-04 | 2.25E+00 | 2.25E+00 |
| HAC4 | AT1G55970 | 1.45E+00 | 9.84E-01 | 9.78E-01 | -1.02E+00 |
| HAC5 | AT3G12980 | 6.24E+00 | 3.78E-01 | 8.99E-01 | -1.11E+00 |
| HAC12 | AT1G16710 | 7.25E+00 | 5.38E-01 | 9.37E-01 | -1.07E+00 |
| HAF1 | AT1G32750 | 1.01E+01 | 5.33E-01 | 1.03E+00 | 1.03E+00 |
| HAF2 | AT3G19040 | 6.40E-01 | 6.91E-01 | 1.37E+00 | 1.37E+00 |
| HAG1 | AT3G54610 | 6.16E-01 | 1.13E-01 | 1.44E+00 | 1.44E+00 |
| HAG2 | AT5G56740 | 1.46E+00 | 8.46E-01 | 9.74E-01 | -1.03E+00 |
| HAG3 | AT5G50320 | 2.15E+00 | 9.48E-01 | 1.01E+00 | 1.01E+00 |
| HAG4 | AT5G64610 | 2.31E+00 | 3.37E-01 | 1.04E+00 | 1.04E+00 |
| HAG5 | AT5G09740 | 9.90E-01 | 8.52E-01 | 1.01E+00 | 1.01E+00 |
| **HMT - Histone methyltransferases** |  |  |  |  |  |
| CLF | AT2G23380 | na | na | na | na |
| SWN | AT4G02020 | 4.41E+00 | 6.76E-01 | 1.06E+00 | 1.06E+00 |
| MEA | AT1G02580 | 2.49E-01 | 3.12E-01 | 1.68E+00 | 1.68E+00 |
| ASHH1/<br>SDG28 | AT1G76710 | 8.92E-01 | 7.30E-01 | 9.14E-01 | -1.09E+00 |
| ASHH2/<br>SDG8 | AT1G77300 | 9.83E+00 | 7.30E-01 | 1.05E+00 | 1.05E+00 |
| ASHH3 | AT2G44150 | 1.05E+00 | 9.66E-01 | 9.80E-01 | -1.02E+00 |
| ASHH4 | AT3G59960 | 1.36E-01 | 3.66E-01 | 1.07E+00 | 1.07E+00 |
| ASHR3/<br>SDG4 | AT4G30860 | 7.48E-01 | 1.69E-02 | 1.74E+00 | 1.74E+00 |
| ATX1 | AT2G31650 | 1.38E+00 | 6.82E-01 | 9.13E-01 | -1.10E+00 |
| ATX2 | AT1G05830 | 3.95E+00 | 3.65E-01 | 8.97E-01 | -1.11E+00 |
| ATX3 | AT3G61740 | 1.50E+00 | 1.12E-01 | 1.37E+00 | 1.37E+00 |
| ATX4 | AT4G27910 | 2.09E+00 | 1.90E-01 | 1.26E+00 | 1.26E+00 |
| ATX5 | AT5G53430 | 2.30E+00 | 3.77E-01 | 8.18E-01 | -1.22E+00 |
| ATXR3 | AT4G15180 | 1.30E+01 | 2.51E-01 | 9.10E-01 | -1.10E+00 |
| ATXR7 | AT5G42400 | 5.34E+00 | 4.30E-01 | 1.09E+00 | 1.09E+00 |
| ATXR5 | AT5G09790 | 2.83E-01 | 5.80E-01 | 1.21E+00 | 1.21E+00 |
| ATXR6 | AT5G24330 | 2.12E-01 | 3.72E-01 | 1.56E+00 | 1.56E+00 |
| SUVH1 | AT5G04940 | 2.28E+00 | 8.73E-01 | 1.01E+00 | 1.01E+00 |
| SUVH2 | AT2G33290 | 1.45E+00 | 4.39E-02 | 1.58E+00 | 1.58E+00 |
| SUVH3 | AT1G73100 | 2.22E+00 | 9.09E-01 | 1.01E+00 | 1.01E+00 |
| SUVH4 | AT5G13960 | 8.94E-01 | 9.20E-02 | 1.54E+00 | 1.54E+00 |
| SUVH5 | AT2G35160 | 1.05E+00 | 5.64E-01 | 1.22E+00 | 1.22E+00 |
| SUVH6 | AT2G22740 | na |  |  |  |
| SUVH7 | AT1G17770 | 2.53E-01 | 3.59E-01 | 2.00E+00 | 2.00E+00 |
| SUVH8 | AT2G24740 | na |  |  |  |
| SUVH9 | AT4G13460 | 4.29E+00 | 6.34E-01 | 9.75E-01 | -1.03E+00 |
| SUVR1 | AT1G04050 | 7.16E-01 | 6.49E-04 | 3.15E+00 | 3.15E+00 |
| SUVR5 | AT2G23740 | na |  |  |  |
| SUVR2 | AT5G43990 | 1.12E+00 | 7.54E-01 | 9.31E-01 | -1.07E+00 |
| SUVR3 | AT3G03750 | 7.91E-01 | 5.08E-01 | 7.98E-01 | -1.25E+00 |
| SUVR4 | AT3G04380 | 6.30E-01 | 3.96E-02 | 1.94E+00 | 1.94E+00 |
| **H3K27me3 demethylases** |  |  |  |  |  |
| ELF6/JMJ11 | AT5G04240 | 5.44E+00 | 6.49E-01 | 9.59E-01 | -1.04E+00 |
| REF6/JMJ12 | AT3G48430 | 6.18E+00 | 6.97E-01 | 1.06E+00 | 1.06E+00 |
| JMJ13 | AT5G46910 | 1.29E+00 | 4.13E-01 | 1.24E+00 | 1.24E+00 |

